# Sequence-intrinsic barriers define a class of translation-restricted uORFs in the human genome

**DOI:** 10.64898/2026.08.12.739961

**Authors:** Hinata Hashimoto, Taichi Akase, Hikaru Kurasawa, Shohei Kitano, Yasunori Aizawa

## Abstract

Ribosome profiling has revealed widespread translation of upstream open reading frames (uORFs) within the human 5′ untranslated region, greatly expanding the apparent coding potential of the genome. However, despite pervasive translational activity, uORF-encoded proteins (uORFps) are only rarely detected by proteomic approaches, creating a major unresolved discrepancy between translation and protein accumulation. Here, we systematically quantified the intracellular accumulation of 111 evolutionarily conserved human uORFps using a unified reporter platform. Although we found that most conserved uORFps accumulated at extremely low levels despite robust transcript expression, protein length emerged as the strongest baseline predictor of this low accumulation. Intriguingly, a subset of conserved uORFs remained poorly accumulated even after experimental extension with a C-terminal EGFP tag, defining a distinct class of translation-restricted uORFs. Sequence and structural analysis of these translation-restricted uORFs revealed a cooperative enrichment of basic amino acids and stable local RNA secondary structures. Synonymous substitutions designed to disrupt these RNA structures substantially restored protein accumulation, demonstrating a major contribution of local transcript stratification to this translation-restricted phenotype. Using these identified rules, we developed a multivariable predictive model that classified 1,432 translation-restricted uORFs across the human transcriptome. Genes harboring these predicted translation-restricted uORFs exhibited significantly reduced downstream translation efficiency and elevated sequence conservation, demonstrating that these sequence-encoded barriers are under purifying selection to act as cis-regulatory elements scattered across the human transcriptome. Together, our findings identify a conserved class of human uORFs that encode intrinsic barriers to productive translation and provide a rigorous framework for understanding how noncanonical coding sequences shape the human proteome.

## Introduction

Upstream open reading frames (uORFs) are short open reading frames located within the 5′ untranslated region (5′UTR) of mRNAs, and approximately half of human transcripts contain at least one uORF^1^. uORFs have long been recognized as cis-regulatory elements that modulate the translation of downstream main open reading frames (main ORFs)^2,3^. Although eukaryotic translation has traditionally been thought to occur preferentially at annotated main ORFs^4,5^, ribosome profiling (Ribo-seq) has revealed widespread ribosome occupancy and translation initiation on uORFs^6–9^.

In addition to regulating downstream main ORFs, some translated uORFs generate protein products, referred to here as uORF-encoded proteins (uORFps)^10–12^. Several uORFps have been detected endogenously and shown to possess biological functions^13,14^. For example, *MIEF1* uORFp localizes to mitochondria, where it regulates mitochondrial translation by binding to the mitoribosome^15^, whereas the *Lin28b* uORFp functions as a reprogramming factor during pluripotency transitions^16^. These examples established that 5′UTR translation can expand the coding potential of human transcripts^7–9^. Yet, such cases remain rare relative to the large number of translated uORFs inferred from ribosome profiling^11,17^.

Despite widespread evidence of uORF translation, the vast majority of predicted uORFps remain undetected by proteomic approaches^11,17^. Mass spectrometry studies typically identify only tens to a few hundred per study, and most reported detections appear in only a single dataset^17–19^. Direct detection of endogenous uORFps by immunoblotting also remains rare^16,20–24^. Consequently, the striking discrepancy between pervasive translational activity inferred from Ribo-seq and the paucity of protein-level evidence highlights a major unresolved issue in uORF research^17,25–27^.

Several mechanisms may explain this long-standing discrepancy between translation and protein accumulation^28–31^. Ribosome occupancy detected by Ribo-seq does not necessarily indicate productive synthesis of stable protein^32^, and uORF-derived peptides may additionally undergo rapid degradation^33,34^. Whether sequence-intrinsic features within uORFs broadly drive this low accumulation and downstream translational regulation remains unclear.

In this study, to focus on uORFs most likely to have functional relevance, we systematically quantified the intracellular accumulation of a comprehensive set of evolutionarily conserved human uORFps. We used C-terminal FLAG reporters to measure uORFp abundance, proteasome inhibition to assess the contribution of post-translational degradation, and experimental extension with a C-terminal EGFP tag, along with N-terminal fusion configurations, to distinguish size- and stability-related effects on productive translation. We then integrated the reporter-derived uORFp accumulation dataset with protein- and RNA-level sequence features to identify the primary determinants of low uORFp accumulation. This analysis allowed us to distinguish simple size-dependent effects from a distinct class of sequence-intrinsic translation barriers. Based on these sequence features, we developed a multivariable logistic regression model to predict this translation-restricted phenotype from sequence features alone. Finally, we applied this model transcriptome-wide, revealing a global landscape of translation-restricted uORFs that operate as sequence-encoded barriers to downstream main ORF translation. Together, our findings provide a mechanistic paradigm for understanding how these noncanonical coding elements systematically shape the human proteome.

## Results

### Conserved uORF-encoded proteins accumulate at low intracellular levels

To systematically evaluate the intracellular accumulation of uORFps, we screened human uORFs from NCBI Refseq dataset (Release 222). We focused on evolutionary conservation as an indicator of potential functional relevance and identified 111 conserved uORFs (cuORFs), which show high amino acid sequence conservation between human and mouse (Fig. 1a and Supplementary Table 1). We next compared their amino acid lengths with those of all human uORFs and canonical proteins. uORFs were markedly shorter than canonical proteins, with a median length of 46 amino acids (Fig. 1b). The median length of the selected cuORFs was 37 amino acids, consistent with the short length distribution of human uORFs.

**Fig. 1.**
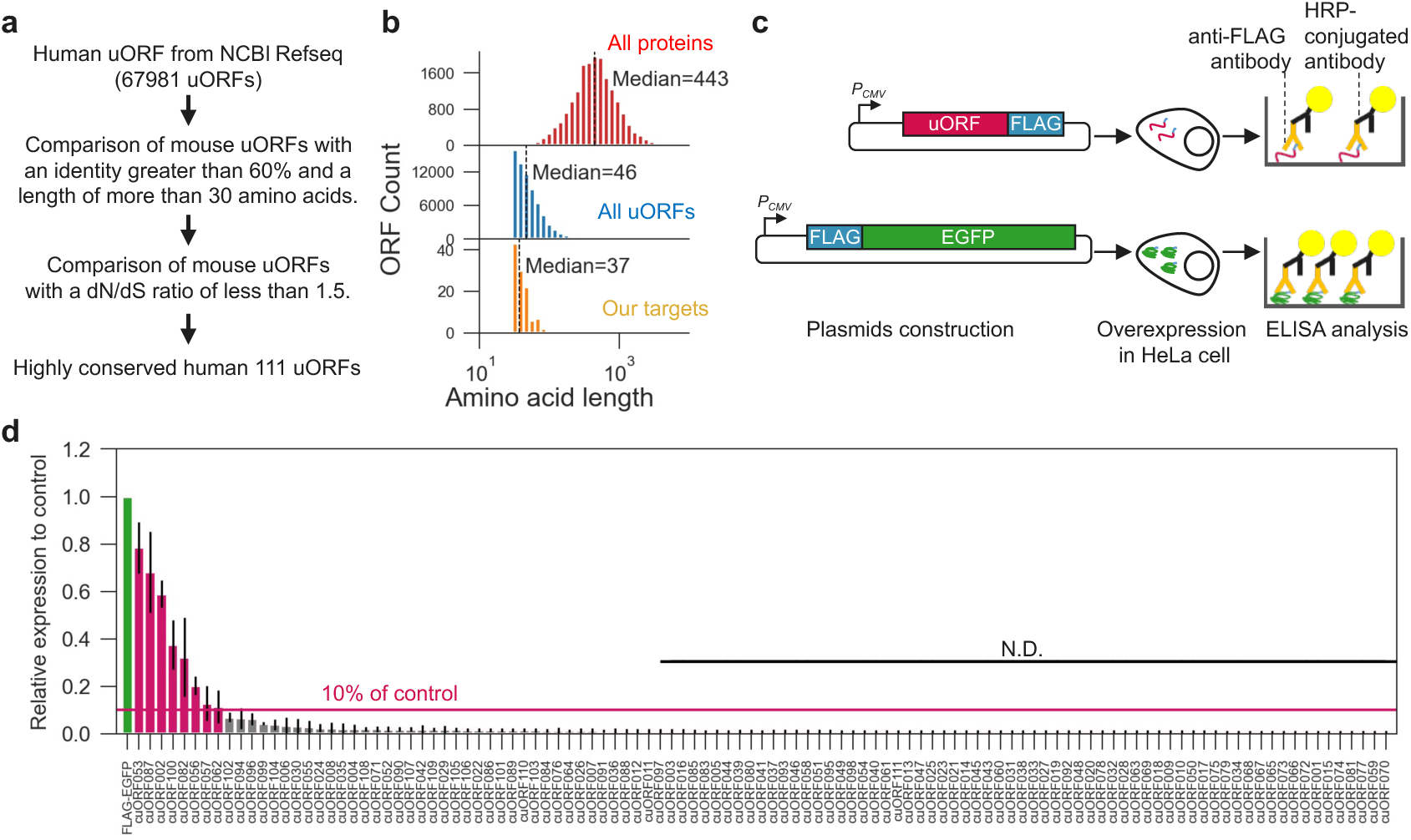
Conserved uORF-encoded proteins accumulate at low intracellular levels. **a**. Schematic overview of the selection of 111 evolutionarily conserved human uORFs from the NCBI RefSeq dataset. Conserved uORFs were defined based on amino acid sequence conservation between human and mouse. **b.** Length distributions of canonical human proteins registered in UniProt (red), all human uORFs (blue), and our targets (orange). **c.** Schematic of the C-terminal FLAG reporter assay used to quantify intracellular accumulation of uORF-encoded proteins. Each conserved uORF was cloned upstream of a C-terminal FLAG tag, and protein abundance was measured by FLAG-based ELISA. **d.** Relative accumulation levels of 111 conserved uORFps measured by FLAG-based ELISA and normalized to the FLAG-EGFP control. Dashed line indicates 10% of the control level.

To quantify intracellular uORFp accumulation, we ectopically expressed each cuORF as a C-terminal FLAG fusion and measured protein abundance using a FLAG tag– based Enzyme-Linked Immunosorbent Assay (ELISA) (Fig. 1c). To ensure a consistent and high level of translation initiation across all constructs, the three nucleotides preceding the start codon were fixed to ACC, providing an optimal Kozak sequence. As a positive control, FLAG-tagged EGFP was used to confirm the sensitivity of the assay. Among the 111 cuORF-encoded proteins (cuORFps) examined, only eight accumulated at levels exceeding 10% of the control (Fig. 1d). Notably, four of these detectably accumulated cuORFps had previously been validated by western blotting or mass spectrometry and were suggested to have cellular functions in prior studies (Supplementary Table 2). In contrast, the remaining 103 cuORFps accumulated at levels below 10% of the control or were below the detection limit. RT–qPCR analysis confirmed that all tested cuORFs were transcribed at levels of at least 10% of the control, indicating that low protein abundance could not be attributed to insufficient mRNA expression (Supplementary Fig. 1a). These results indicate that most conserved uORFps show markedly low intracellular accumulation despite detectable transcript expression.

### Short amino acid length contributes to low uORFp accumulation

To investigate the determinants of low cuORFp accumulation, we extracted sequence features from both high-accumulation cuORFps (>10% of control, n = 8) and low-accumulation cuORFps (≤10% of control, n = 103). We calculated protein-level features including amino acid composition (basic, acidic, and hydrophobic residues), sequence length, amino acid identity with mouse orthologs, and structural confidence based on AlphaFold2 predictions (pLDDT score)^35,36^. RNA-level features included GC content, codon adaptation index (CAI), the dN/dS ratio with mouse orthologs, RNA minimum free energy, and base-pairing probability within the 5′-terminal 15 bp region (Supplementary Fig. 1b, Fig. 2a). Comparison of sequence features between the two groups revealed that low-accumulation cuORFps were significantly shorter than high-accumulation cuORFps (Fig. 2a). The transition between the two groups occurred around 60 amino acids in length.

**Fig. 2.**
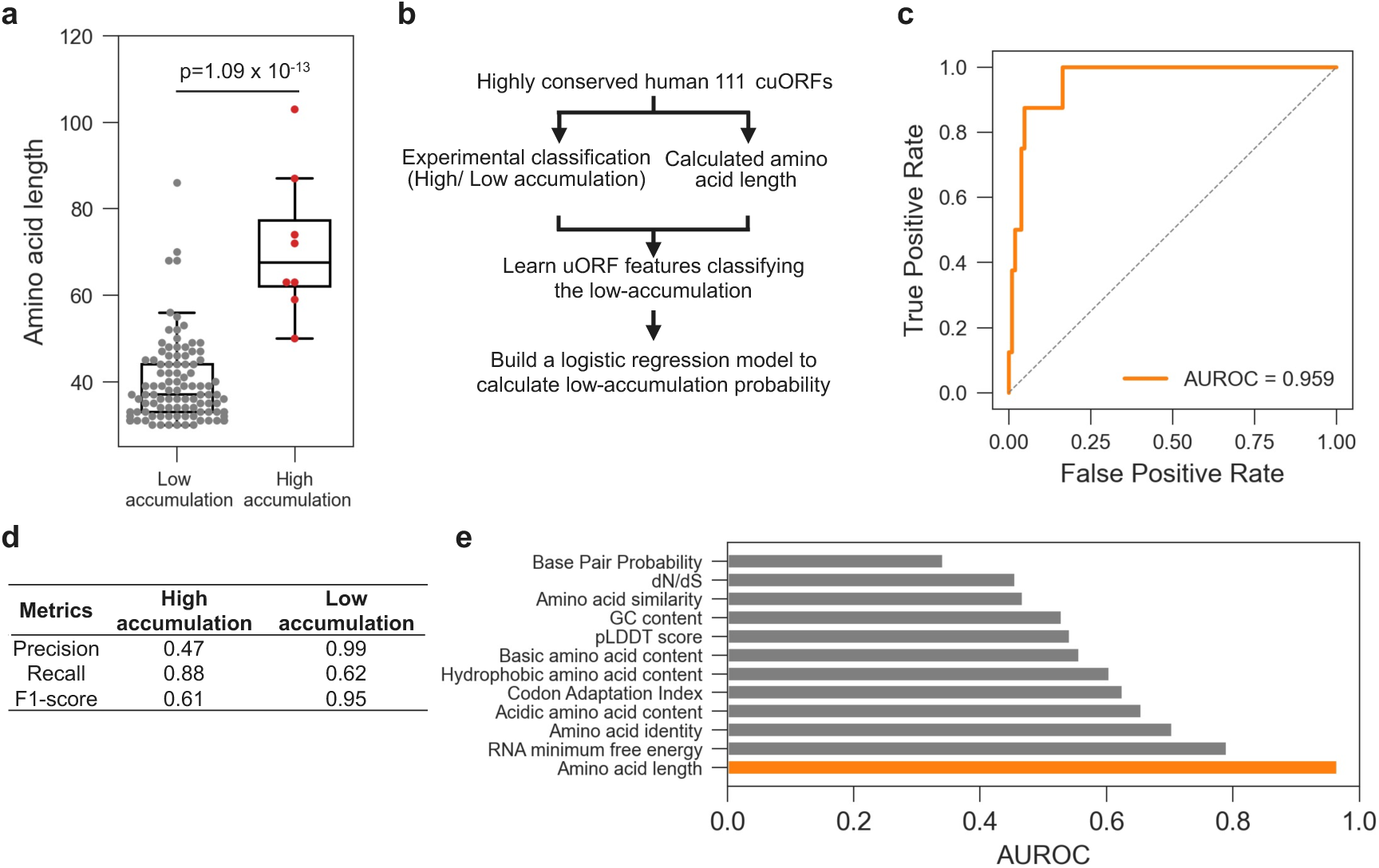
Short amino acid length contributes to low uORFp accumulation. **a.** Comparison of amino acid length between high- and low-accumulation conserved uORFps. **b**. Overview of the logistic regression model used to classify conserved uORFs into high- and low-accumulation groups based on sequence features. The model was trained using 111 conserved uORF reporter constructs. **c**. Receiver operating characteristic curve showing model performance in predicting low-accumulation uORFs. AUROC is indicated. **d**. Confusion matrix showing classification performance for high- and low-accumulation uORFs. **e**. Relative feature importance of logistic regression model.

To determine whether low-accumulation cuORFps could be predicted from sequence features, we next constructed a logistic regression model using the 111 cuORFs. The model was designed to output a probability (hereafter referred to as the low-accumulation probability score) indicating whether a cuORF belonged to the low-accumulation group, which was defined as exhibiting ≤10% accumulation in the cuORFp– FLAG assay. This prediction was based solely on sequence-derived features (Fig. 2b). To minimize overfitting, model performance was evaluated using five-fold cross-validation.

When evaluating individual sequence features, we found that amino acid length alone achieved a high predictive performance, with an AUROC of 0.959 (Fig. 2c). Despite the substantial class imbalance in the training dataset (8 high-accumulation and 103 low-accumulation uORFs), this length-based model retained robust discriminative power for both classes, achieving a recall of 0.88 for high-accumulation uORFs and a precision of 0.99 for low-accumulation uORFs (Fig. 2d). Moreover, this performance by amino acid length was the highest among all sequence features tested (Fig. 2e). These results suggest that protein length is a major determinant of low cuORFp accumulation, although it does not fully explain the behavior of all cuORFs.

### Identification of translation-restricted uORFs

To experimentally evaluate the contribution of protein length to uORFp accumulation, we fused EGFP to the C terminus of each cuORFp-FLAG and quantified reporter accumulation using a FLAG-based ELISA (Fig. 3a). Among the conserved uORFs examined, only 26 exhibited accumulation levels exceeding 50% of the control, whereas 85 showed levels below 50%, including 37 with less than 10% accumulation. These results indicate that increasing protein length can enhance accumulation for a subset of cuORFps. However, many cuORFps remained poorly accumulated despite C-terminal EGFP fusion, suggesting that factors other than protein length also contribute to their low abundance. We therefore focused on this residual group to identify additional determinants of low accumulation.

**Fig. 3.**
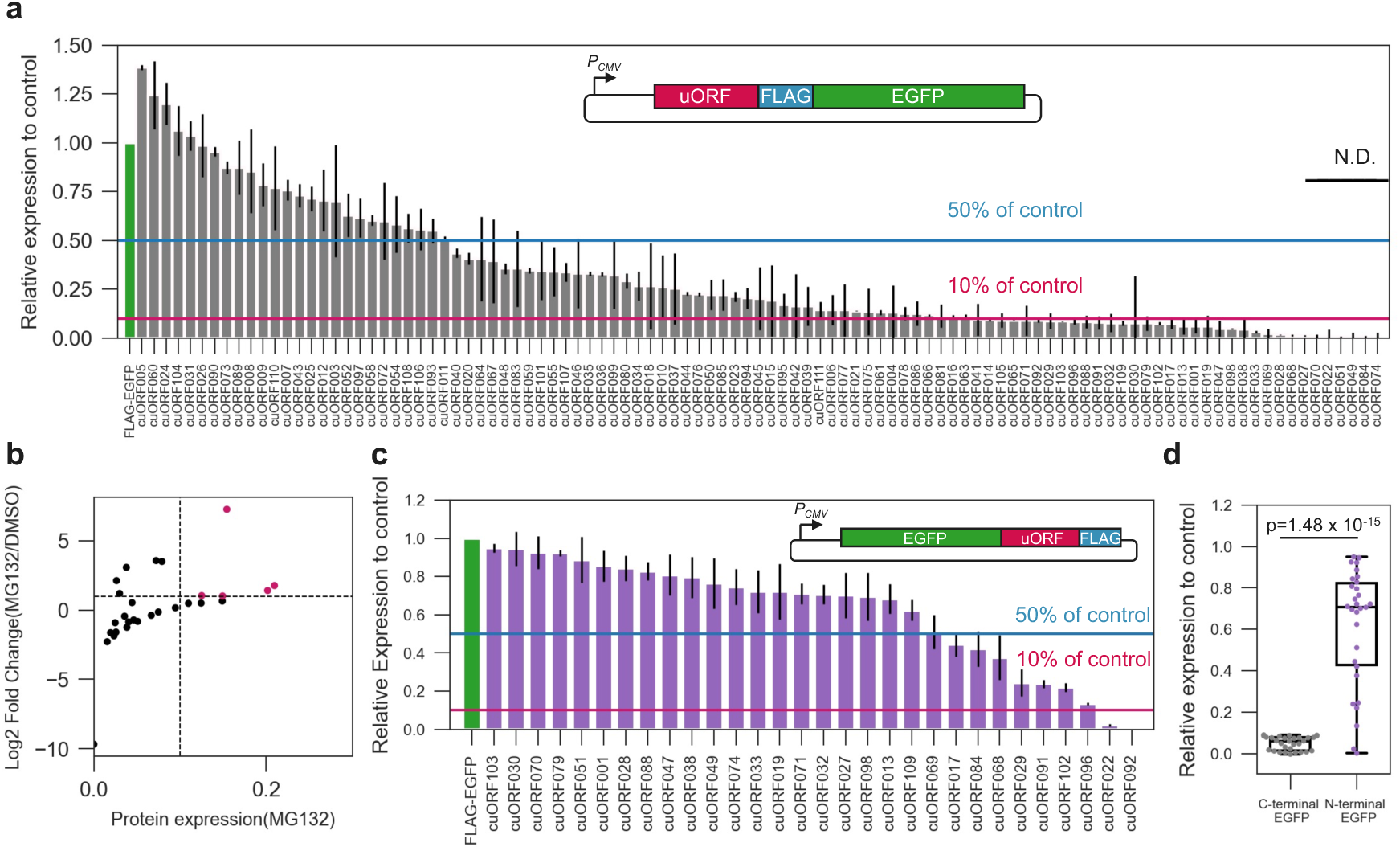
Identification of translation-restricted uORFs. **a.** Relative accumulation of conserved uORFps fused to C-terminal EGFP-FLAG, measured by FLAG-based ELISA and normalized to the EGFP-FLAG control. Dashed lines indicate 10% and 50% of the control level. **b.** Relative accumulation of 30 low-accumulation uORFps after treatment with the proteasome inhibitor MG132. Values were normalized to the EGFP-FLAG control. **c.** Relative accumulation of 30 low-accumulation uORFps fused to N-terminal EGFP, measured by FLAG-based ELISA and normalized to the EGFP-FLAG control. Dashed lines indicate 10% and 50% of the control level. **d.** Comparison of the relative accumulation of uORFp–FLAG-EGFP and EGFP–uORFp–FLAG fusion proteins for 30 low-accumulation uORFps.

One possible explanation for this low accumulation is rapid degradation via the ubiquitin–proteasome system. To test this, we focused on the 30 cuORFps with the lowest accumulation levels, whose accumulation was not restored upon C-terminal EGFP fusion, and quantified their intracellular abundance following treatment with the proteasome inhibitor MG132 (Fig. 3b). Only five of the 30 cuORFps showed partial recovery upon MG132 treatment, with maximal accumulation reaching approximately 20% of the control. These results indicate that proteasome-mediated degradation is not the primary determinant of this low-accumulation phenotype.

Given this, it became likely that an inherent “translation-repressive” property is encoded within the uORF amino acid sequence. We next examined whether this translational restriction depends on the position of the sequence within the full-length proteins by relocating the uORF portions from the N-terminus to the C-terminus of the EGFP module (Fig. 3c, d). In contrast to the uORFp-FLAG-EGFP, 28 of the 30 EGFP-uORFp-FLAG reporters accumulated at levels exceeding 10%, with 21 even surpassing 50%. These findings demonstrate that the position of the uORF within the fusion protein— specifically its location at the N-terminus—is a critical determinant for limiting intracellular accumulation.

This position-dependent restriction implies that the inherent “translation-repressive” property of the uORF is exerted during either translation initiation or the early phase of translation elongation. However, because these constructs utilized an optimal Kozak sequence upstream of the start codon, a deficiency in initiation efficiency is unlikely to account for the low accumulation. Instead, our results demonstrate that within this reporter system, the low accumulation of these uORFps is driven by sequence-dependent mechanisms during early translation elongation. Although further validation is required to determine how these findings translate to endogenous, un-fused uORFs, these results strongly imply that constraints during the early stages of translation play a critical role in the repressive functions of these conserved uORFs.

### Basic amino-acid enrichment and local RNA secondary structure are associated with translation-restricted uORFs

Next, we sought to identify sequence features within the cuORF regions that are associated with early translation resistance. Based on the cuORF–FLAG–EGFP reporter assay (Fig. 3a), cuORFs exhibiting accumulation levels below 10% of the control were classified as translation-restricted uORFs (n = 34), whereas those exhibiting accumulation levels of at least 10% were classified as non-restricted uORFs (n = 69). We then examined the potential RNA and protein sequence features characteristic of this translation-restricted group.

Comparison between the translation-restricted uORFs and non-restricted uORFs revealed significant differences in three features (Fig. 4a-c, Supplementary Fig. 2a). Notably, the proportion of basic amino acids was significantly higher in the translation-restricted uORFs group than in the non-restricted uORFs group (Fig. 4a). This enrichment remained significant when compared with the N-terminal 30 amino acids of all human proteins registered in UniProt^37^. As an extreme example, deletion of the poly-arginine stretch in the *FZD1* uORF restored protein accumulation to the level comparable to the control (Supplementary Fig. 2b).

**Fig. 4.**
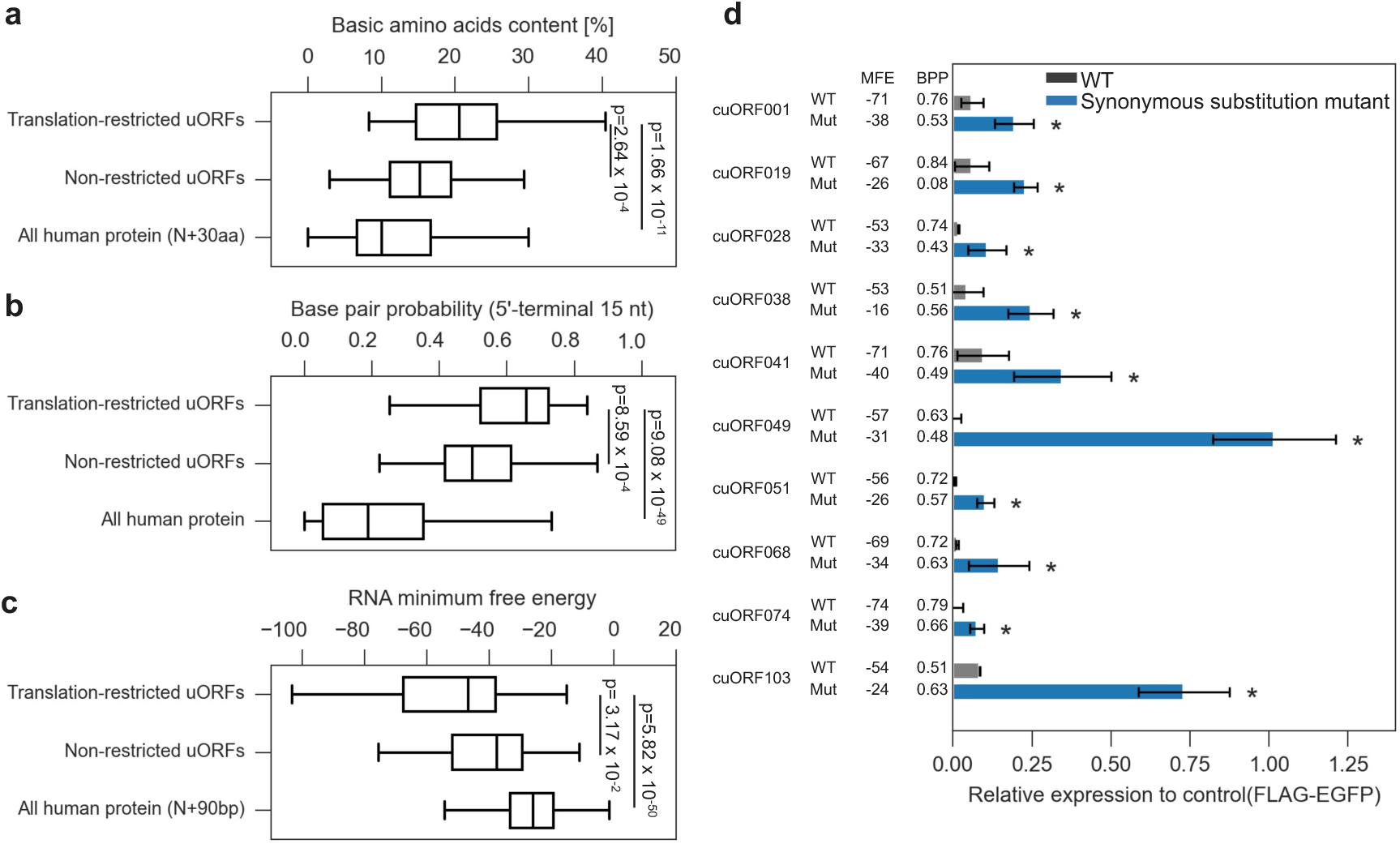
Basic amino-acid enrichment and local RNA secondary structure are associated with translation-restricted uORFs. **a.** Comparison of basic amino-acid content between experimentally defined translation-restricted and non-restricted uORFs, and the N-terminal 30 amino acids of canonical human proteins. **b.** Comparison of base-pairing probability within the 5′-terminal region between translation-restricted uORFs, non-restricted uORFs, and canonical human coding sequences. **c.** Comparison of RNA minimum free energy in the 5′-terminal region between translation-restricted uORFs, non-restricted uORFs, and canonical human coding sequences. **d.** Relative accumulation of synonymous substitution mutants generated for 10 translation-restricted uORFs (*p<0.05). Values were normalized to the FLAG-EGFP construct. MFE denotes minimum free energy, and BPP denotes the average base-pair probability across the 5′-terminal 15 nucleotides of the uORF.

The base-pairing probability within the 5′-terminal 15 bp region was significantly higher in the translation-restricted uORFs group than in the non-restricted uORFs group and all human proteins (Fig. 4b). Consistently, the RNA minimum free energy of uORFs was significantly lower in the translation-restricted uORFs group than in both the non-restricted uORFs group and the 5′-terminal 90 bp of all human proteins (Fig. 4c). A lower minimum free energy reflects an increased propensity to form stable secondary RNA structures^38^. In particular, base-pairing near the start codon (typically within 10–30 nucleotides) has been reported to promote ribosome stalling during the early translation phase^39^.

To test whether RNA secondary structure contributes to the translation-restricted phenotype, we generated synonymous substitution mutants designed to attenuate RNA secondary structure by reducing GC content while preserving the encoded amino acid sequence. We selected 10 cuORFs predicted to form the most stable RNA structures among the 34 translation-restricted cuORFs, which were calculated to have an MFE of less than −50 kcal/mol and a base-pair probability greater than 0.5. The protein abundance of these 10 wild-type cuORFs and their synonymous mutants, expressed as uORF-FLAG-EGFP fusions, was measured by ELISA (Fig. 4d, Supplementary Table 3).

The impact of these substitutions varied substantially among the tested cuORFs. Nevertheless, all synonymous mutants exhibited a clear increase in uORFp accumulation relative to their wild-type counterparts, with several mutants recovering dramatically to levels exceeding 50%. This robust rescue strongly demonstrates that RNA secondary structure is a primary determinant driving the low accumulation of these specific uORFs. Conversely, for the uORFs that showed a more modest recovery, mitigating the RNA structure alone was insufficient to fully counteract the translation restriction. This incomplete rescue indicates that stable RNA structures and basic amino acid enrichment serve as cooperative, multi-layered constraints that robustly enforce translation resistance.

### Sequence features predict translation-restricted uORFs

We next constructed a logistic regression model to predict whether a given uORF belongs to the translation-restricted class, yielding an output termed the translation-restricted probability score. The model was trained using the 103 cuORF–FLAG–EGFP constructs, comprising both high- and low-accumulation groups described above, after excluding uORFs showing ≥10% expression in the FLAG-only reporter to focus on sequences whose accumulation was limited in the absence of EGFP fusion (Fig. 5a). To prevent overfitting, model performance was evaluated using five-fold cross-validation. We trained models using several combinations of features and selected the final model based on a subset of three features. The final model incorporated RNA base-pairing probability, basic amino acid content, and acidic amino acid content. This multi-feature model achieved an AUROC of 0.811 for predicting translation-restricted uORFs (Fig. 5b).

**Fig. 5.**
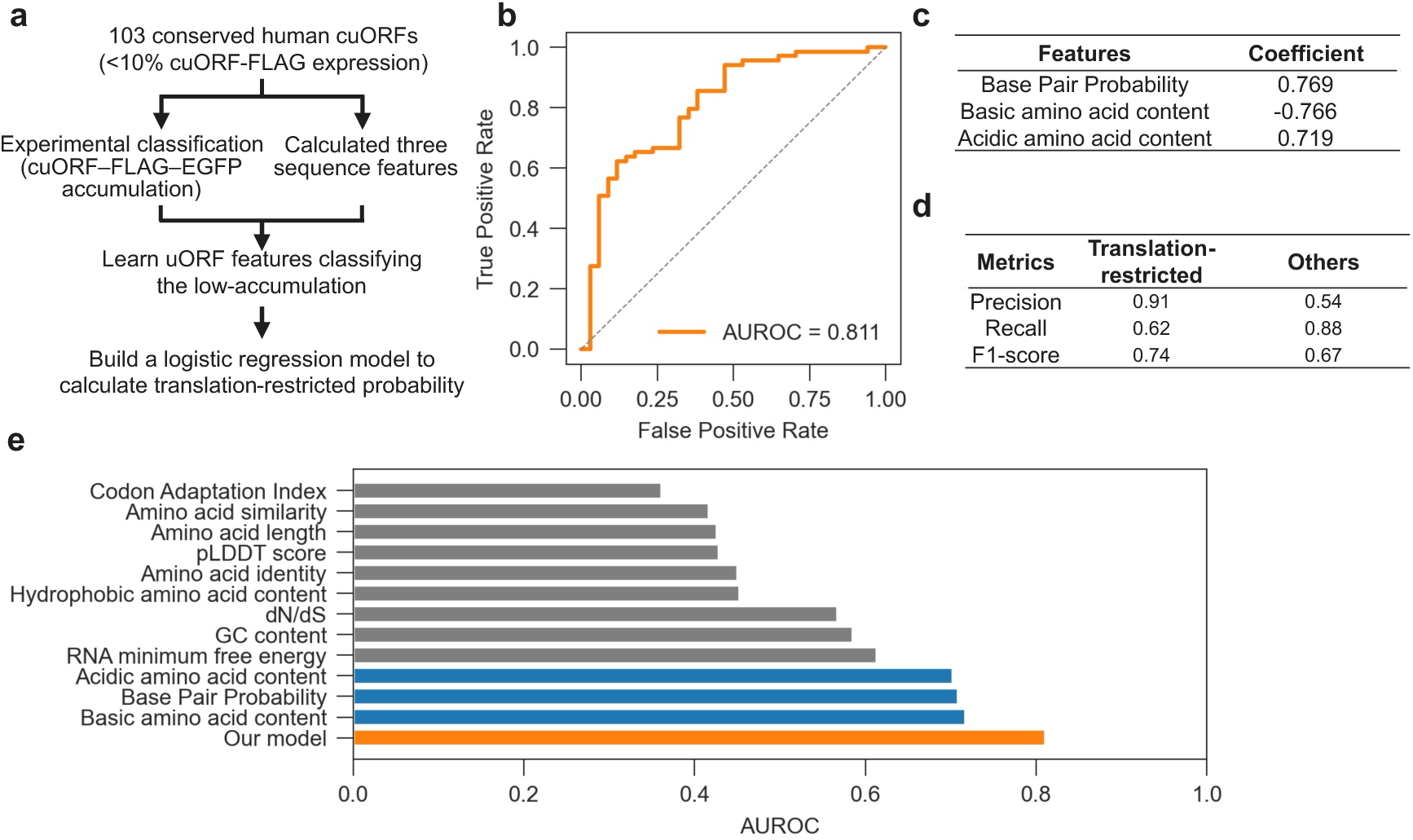
Sequence features predict translation-restricted uORFs. **a.** Overview of the logistic regression model used to classify conserved uORFs into translation-restricted and non–translation-restricted groups based on sequence features. The model was trained using 103 conserved uORF reporter constructs. **b.** Receiver operating characteristic curve showing model performance in predicting translation-restricted uORFs. AUROC is indicated. **c.** Relative feature importance of the final logistic regression model, which incorporated RNA base-pairing probability, basic amino-acid content, and acidic amino-acid content. **d.** Confusion matrix showing classification performance for translation-restricted uORFs. **e.** Comparison of predictive performance among models using individual features or combined sequence features. Blue bars indicate sequence features used as model inputs, whereas orange bar indicate the resulting prediction models.

Feature importance analysis revealed that all three features contributed comparably to the model (Fig. 5c). Notably, the model demonstrated high precision for the translation-restricted class (0.91), albeit with moderate recall (0.62), while showing a complementary trade-off for the non-restricted class with higher recall (0.88) and lower precision (0.54) (Fig. 5d). Incorporating these three features improved classification performance compared with models built on any single feature alone (Fig. 5e). These findings indicate that constraints acting during early stages of translation are unlikely to be explained by a single sequence determinant. Instead, both RNA sequence features and the chemical properties of the encoded nascent protein chains appear to contribute cooperatively to drive the translation-restricted phenotype.

### Transcriptome-wide prediction identifies translation-restricted uORFs associated with reduced downstream translation efficiency and elevated conservation

Finally, we applied the two logistic regression models developed in this study— the low-accumulation predictor and the translation-restricted predictor—to all 67,981 uORFs extracted from the NCBI RefSeq dataset. The low-accumulation model predicted that 60.5% of human uORFs (n = 41,152) belong to the low-accumulation class (Fig. 6a), suggesting that protein accumulation of the majority of uORFs is constrained by their short coding length. Furthermore, the translation-restricted model classified 2.1% of human uORFs (n = 1,432) as translation-restricted (Fig. 6b, Supplementary Table 4). To test whether these predicted translation-restricted uORFs are associated with the translation of downstream main ORFs, we utilized a database that integrates translation efficiency of the main ORFs across multiple Ribo-seq datasets^40^ to compare translation efficiency among all genes (n = 9,194), genes harboring predicted translation-restricted uORFs (n = 590), and genes harboring uORFs with a low predicted translation-restricted probability (n = 4,001) (Fig. 6c). Notably, genes containing predicted translation-restricted uORFs showed significantly lower translation efficiency than the other two groups (p = 8.33 × 10^−6^).

**Fig. 6.**
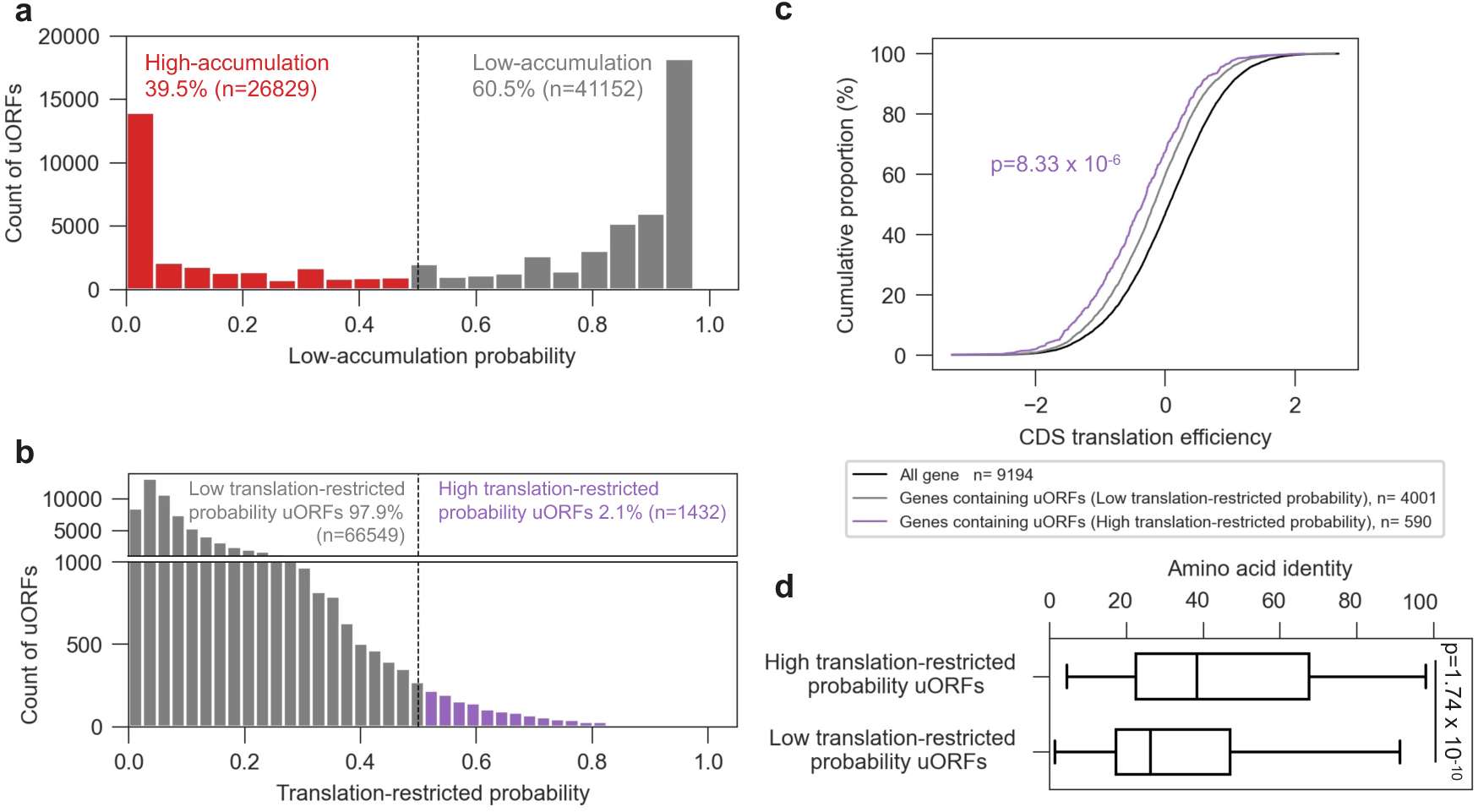
Transcriptome-wide prediction identifies translation-restricted uORFs associated with reduced downstream translation efficiency and elevated conservation. **a.** Transcriptome-wide distribution of low-accumulation probability scores assigned to 67,981 human uORFs from the NCBI RefSeq dataset. uORFs with scores above the defined threshold were classified as predicted low-accumulation uORFs. **b**. Transcriptome-wide distribution of translation-restricted probability scores assigned to 67,981 human uORFs from the NCBI RefSeq dataset. uORFs with scores above the defined threshold were classified as predicted translation-restricted uORFs. **c.** Translation efficiency of downstream main ORFs in all genes, genes harboring predicted translation-restricted uORFs, and genes harboring uORFs with low predicted translation-restricted probability. Translation efficiency values were obtained from an integrated Ribo-seq-based dataset. **d.** Amino acid sequence conservation between human uORFs and mouse orthologous uORFs in predicted high and low predicted translation-restricted groups.

Finally, to examine whether translation-restricted uORFs have been preferentially retained during mammalian evolution, we compared amino acid sequence identity between human uORFs and their mouse orthologs. For this analysis, mouse orthologous uORFs were defined based on amino acid sequence homology to human uORFs, yielding 8,674 human-mouse orthologous uORF pairs. Predicted high translation-restricted uORFs exhibited significantly higher conservation than low translation-restricted uORFs group (p = 1.74 × 10^−10^) (Fig. 6d). Taken together, these results suggest that translation-restricted uORFs define a distinct, evolutionarily conserved subset of human uORFs associated that are preferentially enriched in genes to as repressive cis-regulatory elements that limit downstream main ORF translation efficiency.

## Discussion

This study establishes an integrative, data-driven framework for decoding the human uORFome. First, combining quantitative ELISA data from our cuORF-FLAG reporter assays with systematic sequence feature analyses demonstrated that the low intracellular accumulation of the majority of human uORFps is primarily constrained by their short amino acid length. While this observation is consistent with previous studies suggesting that microprotein stability is influenced by length^34^, systematic experimental evidence linking this parameter to intracellular accumulation across a large set of uORF-encoded proteins has remained limited. By examining a comprehensive set of 111 evolutionarily conserved human uORFps, this study provides a large-scale, empirical dataset directly connecting protein length to uORFp accumulation.

Second, this framework enabled us to discern whether low uORFp accumulation can be explained solely by protein length, leading to the identification of intrinsic translational restriction. While experimentally extending uORFs with C-terminal EGFP fusion constructs substantially influenced accumulation, it did not fully restore the abundance of all cuORFs. This residual low-accumulation phenotype was not broadly rescued by proteasome inhibition using MG132, arguing against rapid ubiquitin-proteasome-mediated degradation as the primary cause. By contrast, placing the EGFP module N-terminal of the uORFp sequence, that is, in a reciprocal configuration, robustly restored protein accumulation. This position-dependent restriction demonstrates that the limiting constraint resides near the uORF N-terminal coding sequence and is exerted during translation initiation or the early phase of translation elongation. As these constructs utilized an optimal Kozak sequence upstream of the start codons, a deficiency in initiation efficiency is less likely to account for this phenotype. Instead, these findings allowed us to propose that the observed restriction operates primarily via sequence-dependent mechanisms during early translation elongation.

Third, this framework capitalized on the unique advantages of our unified reporter configuration to model the sequence-encoded rules of translation restriction. Crucially, while this tag-based expression platform represents an engineered system that does not capture native genomic contexts, it provided a standardized dataset where expression was driven by the same CMV promoter and an upstream optimal Kozak sequence, and detected via RT-qPCR targeting the identical 3’ UTR region and ELISA utilizing an anti-FLAG antibody. This methodological consistency was paramount for filtering out confounding factors, thereby allowing us to isolate and identify the molecular features that drive translation-restricted phenotypes directly from the uORF coding sequences. Our multivariable logistic regression model revealed that translation-restricted uORFs are enriched for basic amino acids and stable local RNA secondary structures, indicating that protein-level and RNA-level properties contribute cooperatively to limit productive translation. The functional relevance of these features was further supported by our synonymous substitution experiments, where disrupting local RNA secondary structure robustly rescued uORFp accumulation. By integrating these precise, sequence-encoded parameters, the resulting translation-restricted predictor achieved high classification performance, establishing a robust computational foundation for genome-wide applications.

Fourth, we expanded this predictive framework transcriptome-wide by applying the two models to all 67,981 human uORFs, translating our assay-driven insights into a global regulatory map. The low-accumulation model predicted that 60.5% of human uORFs belong to the low-accumulation class, reinforcing that short coding length represents a pervasive baseline constraint across the human transcriptome. Concurrently, the multi-feature translation-restricted model identified a subset comprising 2.1% of all human uORFs as translation-restricted. Crucially, transcriptome-wide integration of independent Ribo-seq datasets validated that genes harboring these predicted uORFs exhibited significantly reduced translation efficiency of their downstream coding sequences.

This genome-wide suppression implies that the sequence-encoded early elongation barriers identified in our assays act as widespread, functional bottlenecks for ribosome flux through native 5’ UTRs. This mechanistic constraint conceptually parallels the programmed translational arrest traditionally observed in isolated mammalian paradigms such as the uORFs of *AMD1* or *ATF4*, where specific nascent peptide sequences or local structures slowdown ribosome progression^31,41^. However, while those classical examples have long been viewed as specialized, gene-specific exceptions, our data-driven framework suggests that this translation-repressive architecture operates as a pervasive, system-wide regulatory mechanism across the human genome. Furthermore, the significantly elevated evolutionary conservation of these 2.1% uORFs provides definitive evidence that these translational barriers are not accidental sequence arrangements, but are actively maintained by purifying selection to serve as dedicated cis-regulatory rheostats. Together, these insights propose that translation-restricted uORFs define a distinct, evolutionarily conserved class of functional genomic elements optimized to tune downstream protein synthesis throughout the transcriptome. These findings are complementary to recent large-scale studies reporting that broadly stable microproteins tend to exhibit relatively low evolutionary conservation^10,34^.

Finally, our predictive framework concurrently serves as a screening rationale to prioritize candidate uORFs with an increased likelihood of extended protein half-lives. Although pervasive translation outside of canonical main ORFs is well established^11,12,42,43^, predicting which of these translated products possess the biochemical capacity to accumulate as stable cellular peptides, rather than being rapidly degraded, would highly benefit the field of proteogenomics. Our transcriptome-wide application categorized a substantial population of 26,453 uORFs (approximately 38.9% of the human uORFome) as sequences that conceptually escape both the baseline degradation constraints of short coding length and the early translational barriers identified in this study. This specific subpopulation represents a prioritized pool of candidate uORFs characterized by an increased potential for robust intracellular accumulation and, consequently, the capacity to potentially exert intracellular functions. By offering a sequence-based logic to differentiate translated sequences based on their predicted accumulation phenotypes, these insights complement current mass spectrometry-based proteomic stratification strategies and provide a foundational resource for the future biochemical and functional annotation of stable uORF-encoded microproteins.

Moving forward, a critical next step is to define the precise molecular mechanisms by which these sequence-encoded barriers operate at ribosome-level resolution. Directed approaches such as toeprinting^44^, disome profiling^45^, or single-molecule translation assays^46^ will be indispensable to determine whether individual translation-restricted uORFs act through initiation blockade, elongation slowdown, or ribosome collision. Furthermore, endogenous genome editing of selected uORFs—including targeted disruptions of start codons, basic amino-acid tracts, or structure-forming synonymous sites—will allow us to directly test whether the specific sequence features identified here control both uORFp accumulation and downstream main ORF translation in their native 5’ UTR contexts. In parallel, the molecular mechanisms responsible for the preferential depletion of short uORFps remain to be elucidated. Although previous studies have implicated BAG6-mediated proteasomal degradation in the quality control of short nascent polypeptides^33^, the extent to which this or other length-dependent quality-control pathways contribute to the widespread low accumulation of uORFps remains unknown. This rigorous integration of structural, mechanistic, and genetic approaches will be essential to fully chart the functional landscape of the sequence-based classification proposed in this study.

Ultimately, the sequence-based principles unveiled in this study offer a predictive framework that intersects genomic medicine, synthetic biology, and evolutionary genomics. The ability to predict uORF sequences that constrain productive translation could inform the clinical interpretation of disease-associated 5’ UTR variants^47,48^. This same predictive principle will also be useful for engineering synthetic 5’ UTRs in mRNA therapeutics, where controlled tuning of translation efficiency is essential for the safety and efficacy of genetic medicines^18,49^. Extending this engineering potential to complex architecture, these predictable translation-repressive and accumulation rules can be applied to design eukaryotic artificial operons, offering a programmable mechanism to precisely balance expression ratios among multiple distinct proteins from a single, polycistronic mRNA. Finally, from an evolutionary perspective, the systematic decoding of sequence-intrinsic uORF barriers provides a crucial framework to explain how noncanonical short coding sequences in 5′UTRs are conserved, repurposed, and integrated into mammalian gene-regulatory networks. Beyond uncovering a fundamental layer of post-transcriptional control, this study bridges data-driven genomics with translational science, redefining our understanding of how the dark matter genome shapes the human proteome.

## Methods

### Evolutionary conservation analysis of uORFs

Genome-wide identification and conservation analysis of uORFs were performed as follows. Protein-coding transcripts from the human (hg38) and mouse (mm10) genomes were obtained from Ref-seq datasets. Transcripts containing annotated coding sequences were selected, and 5′ UTRs were extracted. uORFs were identified from the 5′ UTR sequences, and those with lengths between 93 and 453 nucleotides were retained. The corresponding amino acid sequences were generated, and redundant sequences were removed to obtain a non-redundant uORFp dataset. To assess evolutionary conservation, human uORFp sequences were aligned against mouse uORFp sequences using BLAT to identify candidate homologous pairs. These candidate pairs were further refined by pairwise alignment using the Needleman–Wunsch algorithm, and sequence identity and similarity were calculated. uORFs with an amino acid identity of ≥60% between human and mouse and a dN/dS ratio of <1.5 were defined as evolutionarily conserved and used for downstream analyses.

### Plasmid Construction

ELISA analyses plasmids were constructed using the following protocol. First, DNA fragments encoding the CMV promoter, EGFP and FLAG tag were amplified by PCR using KOD One PCR Master Mix (Toyobo). Then, 5′ UTRs were PCR-cloned from cDNAs prepared with Human Total RNA Master Panel (Clontech) and SuperScript IV Reverse Transcriptase (Thermo Fisher Scientific) into the pcDNA3 using NEBuilder HiFi DNA Assembly (New England Biolabs). All primers used for plasmid construction are listed in Supplementary Table 5.

For the construction of synonymous substitution mutants, DNA fragments were automatically designed using a custom script, amplified by PCR, and assembled into pcDNA3 using the NEBuilder HiFi DNA Assembly Kit (New England Biolabs). All DNA fragments used for plasmid construction are listed in Supplementary Table 6.

### DNA sequencing

DNA sequencing was performed using the BigDye Terminator v3.1 Cycle Sequencing Kit (Thermo Fisher Scientific) on a 3730xl DNA Analyzer (Applied Biosystems).

### Cell culture and transfection

HeLa cells were maintained in DMEM (4.5g/l Glucose) (Nacalai Tesque) supplemented with 10% FBS. Cell line was cultured in a humidified incubator with 5% CO2 and 95% air at 37 °C. Transfection into HeLa cells was performed using FuGENE HD (Promega), following the manufacturers’ procedures. For the ELISA analysis, 6×10^4^ cells were cultured in 96-well plates.

### RNA Purification and Quantitative RT-PCR

Total RNA was extracted using RNeasy Mini Kit (Qiagen) and reverse-transcribed by PrimeScript RT reagent Kit with gDNA Eraser (Takara). Real-time PCR was performed using SYBR Green qPCR Master Mix (Thermo Fisher Scientific) and a QuantStudio 6 (Thermo Fisher Scientific) according to the manufacturer’s protocols. 18S rRNA was used as an internal control for normalization. The relative fold-changes were calculated by the ΔΔCt method. All primers used for RT-PCR are listed in Supplementary Table 7.

### ELISA analysis

At 48 h post-transfection, cells were lysed in 5 μL of SDS sample buffer (0.06 M Tris-HCl (pH8.0); 5% Glycerol; 1.7 % SDS) and collected. The lysates were diluted 800-fold with PBST, and 50 μL of the diluted samples were added to MaxiSorp Black plates (Nunc) and incubated at 4°C for 24 h. After removal of the solution, wells were washed with PBST and then blocked with 200 μL of blocking buffer (2% Perfect-Block (MOB) in PBST) at 25°C for 10 min. After washing with PBST, 50 μL of anti-FLAG antibody (Fujifilm), diluted 1:5,000 in blocking buffer, was added to each well and incubated at 25°C for 1 h (primary antibody reaction). Following washing with PBST, 50 μL of HRP-conjugated sheep anti-mouse IgG (GE Healthcare), diluted 1:5,000 in blocking buffer, was added and incubated at 25°C for 1 h (secondary antibody reaction). After washing with PBST and DPBS, enzymatic reactions were performed using the QuantaRed™ Enhanced Chemifluorescent HRP Substrate Kit (Thermo Fisher Scientific) according to the manufacturer’s instructions. Fluorescence was measured using an EnSpire plate reader (PerkinElmer).

### Sequence feature analyses

Base-pairing probability was calculated using the ViennaRNA package^38^. RNA partition functions were computed for each uORF sequence, and base-pair probabilities were obtained from the resulting probability matrix. The mean base-pairing probability of the first 15 nucleotides from the 5′ end was used as the representative Base-pairing probability value for each uORF. Codon adaptation index was calculated based on codon usage frequencies in the human genome^50^. Relative adaptiveness values were assigned to each codon according to synonymous codon frequencies, and the geometric mean of values across all codons in each uORF was calculated. Stop codons were excluded from the calculation. Protein structures of uORFp were predicted using ColabFold, an AlphaFold2-based structure prediction pipeline^51^. Mean predicted local distance difference test (pLDDT) scores were calculated from the predicted models and used as indicators of structural confidence.

### Logistic regression analysis

To identify sequence features associated with uORF behavior, two independent logistic regression models were constructed: one to predict low uORFp accumulation and the other to predict translation-restricted uORFs. For both models, all calculated sequence and structural features were initially extracted as candidate explanatory variables and standardized using z-score normalization prior to model fitting.

For the low-uORFp accumulation model, uORFp accumulation was evaluated using the uORF–FLAG reporter. uORFs with relative expression levels <10% of the FLAG-EGFP control were assigned to the low-accumulation class, whereas those with expression ≥10% were assigned to the high-accumulation class.

For the translation-restricted model, translational repression was evaluated using the uORF–FLAG–EGFP reporter. To exclude uORFs with intrinsically high protein accumulation, sequences showing ≥10% expression in the uORF–FLAG reporter were excluded before model construction. The remaining uORFs were classified according to their FLAG–EGFP reporter expression, with expression <10% defined as the translation-restricted group and expression ≥10% defined as the non-translation-restricted group.

For each model, informative features were selected from the initial candidate feature set based on model performance, and the final feature combinations differed between the two prediction tasks. Logistic regression with L2 regularization was performed using the liblinear solver with class-weight balancing. Model performance was evaluated using stratified five-fold cross-validation, and the area under the receiver operating characteristic curve (AUROC) was used as the evaluation metric. To assess the predictive contribution of individual features, additional models were constructed using selected features or feature combinations and evaluated using the same procedure.

## Acknowledgements

We thank Hitoyoshi Yamashita, Tomoyuki Ohno, Hiroyuki Tsutsumi, Kiyoshi Yamaguchi and the other members of the Aizawa Laboratory for their insightful discussions and invaluable experimental assistance. We also thank Dr. Hideki Taguchi and Dr. Tatsuya Niwa for their helpful comments and discussions. Additionally, we thank the Integrative Bioscience Facility at the Institute of Science Tokyo for providing DNA sequencing services. This work was supported by the Japan Society for the Promotion of Science (JSPS) KAKENHI (Grant Number: 23KJ0939 to H.H., 16K15116 to Y.A., and 16J08765 to S.K.).

## Author contributions

H.H., T.A., H.K., S.K. and Y.A. formulated the concept of this research and designed experiments. H.H. and T.A. performed the experiments. H.H. and T.A. provided the computational analyses. H.H. and Y.A. wrote the manuscript. All authors discussed and commented on the final draft of the manuscript.

## Competing interests

The authors declare no competing interests.

## Notes

### Competing Interest Statement

The authors have declared no competing interest.

