## Supplemental figures and tables for "Sequence-intrinsic barriers define a class of translation-restricted uORFs in the human genome"

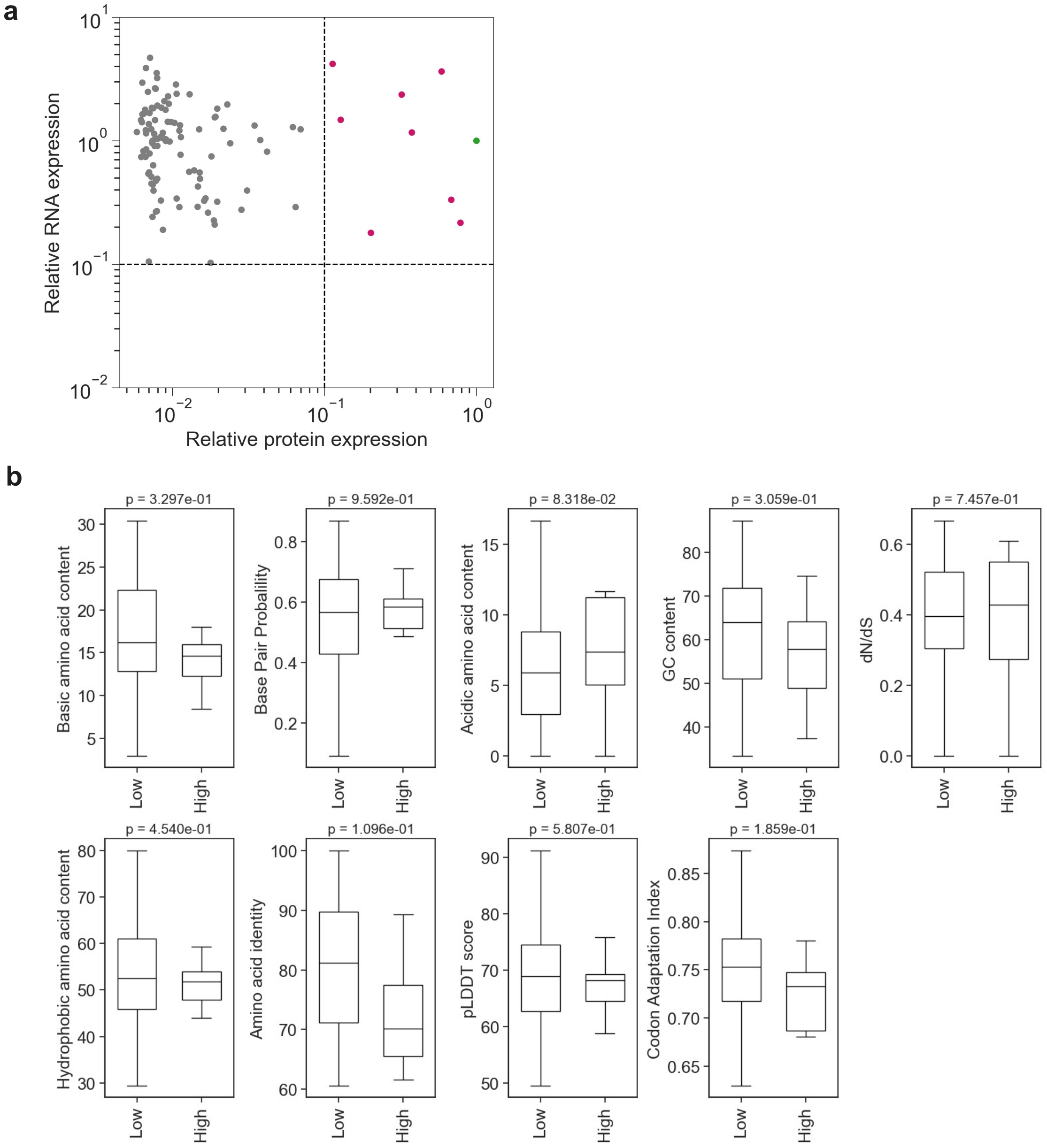


**Supplementary Figure 1 | Characterization of high-accumulation uORFps.** **a**. Relationship between RNA accumulation levels and protein accumulation levels for the 111 cuORFs analyzed in this study. Each dot represents an individual cuORF-FLAG. The green dot indicates the FLAG-EGFP control, whereas red dots represent cuORFs whose protein accumulation levels exceeded 10% of the control. **b.** Comparison of sequence features between low- and high-accumulation conserved uORFps.

**a**

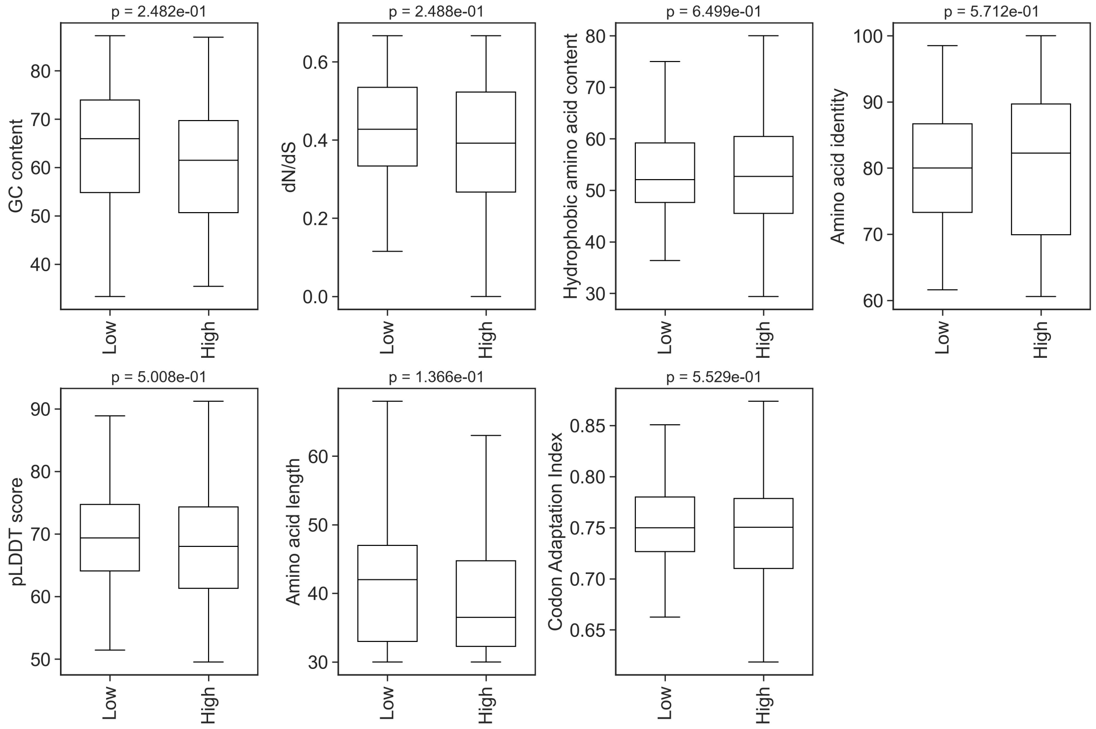


**b**

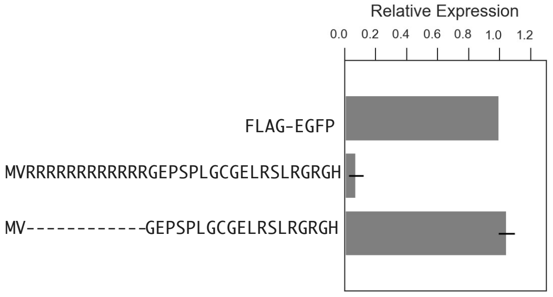


**Supplementary Figure 2 | Sequence features of translation-restricted uORFps.**

**a.** Comparison of sequence features between translation-restricted uORFps (Low) and non-restricted uORFps (High). **b.** Relative accumulation levels of FLAG-EGFP fused to the *FZD1* uORF and of a mutant lacking the N-terminal stretch of consecutive basic amino acids.

### Supplementary Table 1 | cuORF list.

| cuORF ID | Refseq ID | DNA seq | Protein seq | Amino acid identity to mouse homologs (%) |
|---|---|---|---|---|
| cuORF001 | NM_001008701 | ATGTGCCGGGCGGGGGGCCCGGGTTCGCCGAGCCGCAGGAGAGACACGCTGGGCCGACCCCAGAGAGGCGCTGGACAGGCTGGTGGTCCAGGCCGTGGTGCCTGCCAGGTGATGTGGGGCAAAGCCCCCCGCACAGGCCACTGA | MCRAGGPGSPSRRRDTLGRPQRGAGQAGGPGRGACQVMWGKAPRTGH | 66 |
| cuORF002 | NM_001173480 | ATGGACAGTCTGACAGAACAGAGACTGACATCTCCCAATCTGCCGGCCCCCCACCTGGAACACTACAGTGTTCTGCATTGCACCATGACCCTGGATGTGCAAACTGTAGTCGTTTTTGCCGTGATTGTAGTCCTCCTGCTTGTCAATGTCATACTCATGTTTTTCCTGGGAACGCGCTGA | MDSLTEQRLTSPNLPAPHLEHYSVLHCTMTLDVQTVVVFAVIVVLLLVNVILMFFLGTR | 100 |
| cuORF003 | NM_001267617 | ATGGGACTCCAGTTGTCTCTGATCACTTGTGTGGATTTTCCTGGCGTAGAACGACAGAAGCCGCTAGTAAGTCGCCAAGACCTACAGCAGGAATTCTGCACCAAAGGGCATAAAATCTTGTTATTTTAA | MGLQLSLITCVDFPGVERQKPLVSRQDLQQEFCTKGHKILLF | 64.3 |
| cuORF004 | NM_014382 | ATGCTGCTGCTAGGGGTGGTGGGAGCAGCCGTGGGACGCGTGGCCGGGAGCGGGGGTGACAGCCTGGGATTCCGGGGGCTTCTCTTCCTTGTCCTCCTCCTCTCCTCTCTATTCCCAGTGTGGCCGTGGCTGACACTAAAGACTTTGTAG | MLLLGVVGAAVGRVAGSGGDSLGFRGLLFLVLLLSSLFPVWPWLTLKTL | 98 |
| cuORF005 | NM_207303 | ATGCCCGAGTGCGACTGGAGGCAGCCGAGCGGAGGCGACGGCGGTTGGGATCTGTCCCTCCTGACCGGGGAGCGGGACTCGGACGGGCGCCGGTGA | MPECDWRQPSGGDGGWDLSLLTGERDSDGRR | 64.5 |
| cuORF006 | NM_004326 | ATGCGAGATTTTCCTCTGGCAGCAGGAGGCACGCACCCAGAGAATGCTGGAGCTGCAAGGGGAAAGGACCCACTTCCACAGCAGAGAAAAACAAAGAGGAAAAAGGCATACAGGCAGCGAGCGCTAAGGGACGCACCCAGCAAGCAGTGGGCCAGTGCCACTGCCCCCAGCAGCTGTTTCTGCTGCAACCCGAGAGGAACTCGGTGA | MRDFPLAAGGTHPENAGAARGKDPLPQQRKTKRKKAYRQRALRDAPSKQWASATAPSSCFCCNPRGTR | 83.8 |
| cuORF007 | NM_001002860 | ATGGATGAAAAGAGGCTGCTGGATCCAGGGTTGAAGGTTCATAAAGGCCTCTGGGATTATACTCTGAAGAACCCGCAGACGGAATGCCTGAGTGACTGA | MDEKRLLDPGLKVHKGLWDYTLKNPQTECLSD | 84.4 |
| cuORF008 | NM_001082534 | ATGTCTTCTTGGGGCGCCAGACGAATCGGGGTCTCGTTTTTGCTGGAAGAGCCCAGTGTTGGTGGCTTCAGGTGGCTGCTGCCGCCGCCGCCGCCGCCGCCGCTGCTAGTGCGGTTTCCGCCGCTGGTGCGAAGAGAAGAGACACGCGAGCGGGGAGACCTCCAAGGCAGCGAGGCATCGGACATGTGTCAGCACATCTGGGGCGCACATCCGTCGAGCCCGAGGGGAGATTTGCCGGAACAATTCAAACTGCGATATTGA | MSSWGARRIGVSFLLEEPSVGGFRWLLPPPPPPLLVRFPPLVRREETRERGDLQGSEASDMCQHIWGAHPSSPRGDLPEQFKLRY | 75.6 |
| cuORF009 | NM_001017440 | ATGGAGTTTCTAGGGCGCGCAACTCCAGCGGGGCCTCGGCCACTGGGCTCCGGGGCGCAGCGAGGCAGCCGCGCACATGGGCTGAACCCGCGCGGCCAGCGGGAGCGGGAGGGGTAG | MEFLGRATPAGPRPLGSGAQRGSRAHGLNPRGQREREG | 89.7 |
| cuORF010 | NM_020753 | ATGCTGGCTGCTCTCTGGGTGGGCACTGGTGTTCCACAGCCAAGCCGGGAAGATTGGTGTGCCCAGTCCCAGCCCATCCTGGAGCTGTCGGAAGCCACAAGCTGA | MLAALWVGTGVPQPSREDWCAQSQPILELSEATS | 67.6 |
| cuORF011 | NM_001796 | ATGAACGGAATCCGGAGGCGCCTGCCTAGCGAGCGAGGACCGGTCGGCGCTTGCCGCCCTCGGGAGCTAACCCCGAGCACCTCCAGCCATTTGTGA | MNGIRRRLPSERGPVGACRPRELTPSTSSHL | 61.3 |
| cuORF012 | NM_006201 | ATGGCTGAAGACTACTGGGACGGGCGCCTGCGGCGAACAGGAGGAGAAGGAGGTCGCGCGGCCTCATCCCGGGCCGCCGCCCCAGGCGCCGCCGCGCCGGCCCCGCGGCTCTGA | MAEDYWDGRLRRTGGEGGRAASSRAAAPGAAAPAPRL | 94.6 |
| cuORF013 | NM_003948 | ATGCGGCCCGTGGTCCTCCTAAGTGTGAGCTTGCGGCGGACCGAGGCCCACCTGCCTCCCTGCCTGCTTCGCCCTGGACTCGTGACTGCGTCCGCAGAAGAAATCACAACAGCGCTGGAATTGCTAGTTTGCTAG | MRPVVLLSVSLRRTEAHLPPCLLRPGLVTASAEEITTALELLVC | 75 |
| cuORF014 | NM_182553 | ATGAGCGATCGGGGCCCGGGCAGCCGGCAGCGGACGCGCCCCCCGAGCCCACCGGCCCGCGCCCCGCGCCCCCCACGGCCCCGCGGTCCCGGTCCCGGCCGCATCACCCACGTCCCCCGAGCCCCACGGGCCATGCCCGGCCGGCCCTAA | MSDRGPGSRQRTRPPSPPARAPRPPRPRGPGPGRITHVPRAPRAMPGRP | 83.7 |
| cuORF015 | NM_032888 | ATGGGCCTGGCGCGGGCGCCCGCGGGGCCCCAGCCGCGCTGCCTGCCTGCTCGGGCGCCCCTGGGCGCGGGGCTGCGCTGGGGGCGCGGGGGCCGCGCGCTCTAA | MGLARAPAGPQPRCLPARAPLGAGLRWGRGGRAL | 67.6 |
| cuORF016 | NM_030627 | ATGAGGGAAAAAAACCCCGGAGCCGCAGAGAGGGGAAGAGGCAGAAACCGCAGGACCTTCCAGGTCGCCCCCTCGGTCCCCGCACCCCCAGGCCGCCCGCTCACCCTCGTCGAGTCTCGCTAA | MREKNPGAAERGRGRNRRTFQVAPSVPAPPGRPLTLVESR | 87.5 |
| cuORF017 | NM_001012614 | ATGAGGCTCCGAGACAGGACGCGGTTCTGCCTGGGGATCCTGAAGATAAAAAGCTTTGAAAAGTCGAATTCATGGTCGTGGAAGCTGAGCCCATATTAA | MRLRDRTRFCLGILKIKSFEKSNSWSWKLSPY | 75 |
| cuORF018 | NM_001195057 | ATGTTAAAGATGAGCGGGTGGCAGCGACAGAGCCAAAATCAGAGCTGGAACCTGAGGAGAGAGTGTTCAAGAAGGAAGTGTATCTTCATACATCACCACACCTGA | MLKMSGWQRQSQNQSWNLRRECSRRKCIFIHHHT | 94.1 |
| cuORF019 | NM_001243254 | ATGCGGTGCGGGGCGGCCCGGTGCCCCCCCGCCCCGTCACGGCAGCCGCGGCGGCCGAGGGGACCGGGCCAGGGCCGGGGGCGGCGGCCCGAGCCGCGGTAG | MRCGAARCPPAPSRQPRRPRGPGQGRGRRPEPR | 75.8 |
| cuORF020 | NM_001300983 | ATGTCGTTTTGTAGTCAGAGAAAACAGAGATCAATGCATTTTCAAACTGACAGAGGGAACGGATGCTCTTTAGTAGCACATGCCCAGGATCGTGTGTGTGGGGCTTGCGCTGTGCTGAGAAGCTGA | MSFCSQRKQRSMHFQTDRGNGCSLVAHAQDRVCGACAVLRS | 95.1 |
| cuORF021 | NM_001399 | ATGCGGCTGCTCCCTGCTCCGTCCCGCCCAGCCACTGTCGCGCAGGAACGGGTCCCTGCAGCCCCCAGCCGATGGCAGGACAGTAGCCGCCTGTCAGAGGTCGTGAACGGCTGA | MRLLPAPSRPATVAQERVPAAPSRWQDSSRLSEVVNG | 86.5 |
| cuORF022 | NM_080732 | ATGGCGCGCTGTGCGGCGCAGGGCGGCTGGCACAAACGGCGGCGCCGGGGCCGGAGGAAAAAGCTCGCCACCCTGAAGGGTCCCTTCCCAAGCCCTTAG | MARCAAQGGWHKRRRRGRRKKLATLKGPFPSP | 81.2 |
| cuORF023 | NM_201999 | ATGCTTAGACTACTTAACATACAAACTGCTTTCTGGTTAATCATCTTTAGAAGACTGGATTTCTGGATATCTACTCCACTCCATCTCTATTGA | MLRLLNIQTAFWLIIFRRLDFWISTPLHLY | 73.3 |
| cuORF024 | NM_004730 | ATGCAGCTGCCTGGAGAGAGGGAGCCGGTGTCCTACGTCAGAGCCGCCGCCGCCGCGGAGCCGCCGCCGGGGAGGAGCAGCCGCTGCCGCCCAGGACTGGGCCCTTAG | MQLPGEREPVSYVRAAAAAEPPPGRSSRCRPGLGP | 100 |
| cuORF025 | NM_000127 | ATGCAAGAAAATTACAGTCCGCTGCTCGCCCGCCCTGGGTGCGAGATATTCAGCCCCGCTCTCTCCCGTGCATTGTGCAACCCAAAGATGAAAGACCGAAGGGGAGAAAGTTAA | MQENYSPLLARPGCEIFSPALSRALCNPKMKDRRGES | 91.9 |
| cuORF026 | NM_001039762 | ATGCCCTGTCCGGAGTGCTCCGTGCCGCCCCAGGCTTGCCAGGGTCCGCAGCGAGACATGCCCGAGCCAGGCGCTGCATCGACCTCGACTTATCTGTAA | MPCPECSVPPQACQGPQRDMPEPGAASTSTYL | 81.2 |
| cuORF027 | NM_178539 | ATGCCTAGAAGGGCAGCGGGTGGGCTGCGTGGGGCTCTGCCCGCAGCACTTCGAGGCGGGATTGAGGGGCTGAAGCTGGCCAGGAGTTGCTGCTAA | MPRRAAGGLRGALPAALRGGIEGLKLARSCC | 77.4 |
| cuORF028 | NM_025182 | ATGGTGGCAGTCACCTTGCCAGTGCAGCAGGCATGGATCAGACTGCAAGCTAGGGGCACGAGGCATCAGTTGGGAAGAGGGGACGCACAGCTAGGCTTGAGGCCTCTTCATCGGGATGTCCCCAGGCCCCCCAGCCCAGGCCCAGCATAA | MVAVTLPVQQAWIRLQARGTRHQLGRGDAQLGLRPLHRDVPRPPSPGPA | 81.6 |

### (Continued)

| | | | | |
|---|---|---|---|---|
| cuORF029 | NM_001037163 | ATGGCGGCGGCTCTGTCGGGCCTGGCTGTCCGGCTCTCGCGCTCGGCCGCCGCCCGCTCCTATGGGGTCTTCTGCAAGGGGCTGACTCGCACGCTGCTCATCTTCTTCGACCTGGCCTGGCGGCTGCGTATCAACTTCCCCTACCTCTACATCGTGGCTTCCATGATGCTCAACGTCCGCCTGCAGGTTCATATTGAGATCCATTGA | MAAALSGLAVRLSRSAAARSYGVFCKGLTRTLLIFFDLAWRLRINFPYLYIVASMMLNVRLQVHIEIH | 98.5 |
| cuORF030 | NM_005250 | ATGAAGAAGGGACAGAGCACGGAGCGGCCGGGGAGAGCGGCAGAGCCAGAGGCAGAGGCACGGCTGGCTCCCCGGGAGGGCCCTTGCGGCGCGGGGCGCAGTGCCTAG | MKKGQSTERPGRAAEPEAEARLAPREGPCGAGRSA | 62.9 |
| cuORF031 | NM_148898 | ATGCCATCAGTCTGGGACGTGATCGGGCAGAGGTGTACTCACAGTAGTGTAAATACTGCTGTAAATAGTTGTCTGATGGTGGCTTTGACAGTGAGCTAG | MPSVWDVIGQRCTHSSVNTAVNSCLMVALTVS | 100 |
| cuORF032 | NM_003505 | ATGGTCAGGCGGCGGCGGCGGAGAAGGAGGCGGAGGCGCAGGGGGGAGCCGAGCCCGCTGGGCTGCGGAGAGTTGCGCTCTCTACGGGGCCGCGGCCACTAG | MVRRRRRRRRRRRGEPSPLGCGELRSLRGRGH | 70.6 |
| cuORF033 | NM_017412 | ATGTTGGCCAAATGTGCCCCATGTAATAAAATGAAAAGAAGAGACAAGATGATGTCATTTTCCCATATTGTGAAACCAAAAACAAACGCCTTTTGTGAGACCAAGCTAACAAACCTCTGA | MLAKCAPCNKMKRRDKMMSFSHIVKPKTNAFCETKLTNL | 79.5 |
| cuORF034 | NM_006143 | ATGAATGTGTTTCCAGGCTTTCCTGGTGGTTTATGGCATTCTCCAAACTCCTATGCAAGGGCTATTCCTGACCAAGAAGATCTAAAGAGAACGTCTCTGAAATCAAGTCCGGATGAAGAATTAAGAGAAAAAAAGTGA | MNVFPGFPGGLWHSPNSYARAIPDQEDLKRTSLKSSPDEELREKK | 83.3 |
| cuORF035 | NM_000840 | ATGAGGAGGACCAACCATGAGCCAGAGCCCGGGTGCAGGCTCACCGCCGCCGCTGCCACCGCGGTCAGCTCCAGTTCCTGCCAGGAGTTGTCGGTGCGAGGAATTTTGTGA | MRRTNHEPEPGCRLTAAAATAVSSSSCQELSVRGIL | 83.3 |
| cuORF036 | NM_001146156 | ATGTCCCCGGCGAATGGGGAACAGTCGAGGAGCCGCTGCCTGGGGTCTGAAGGGAGCTGCCTCCGCCACCGCCATGGCCGCTGGATCCAGCCGCCGCCTGCAGCTGCTCCTGGCGCAATGAGGAGAGGAGCCGCCGCCACCGCCACCGCCCGCCTCTGA | MSPANGEQSRSRCLGSEGSCLRHRHGRWIQPPPAAAPGAMRRGAAATATARL | 89.7 |
| cuORF037 | NM_001113239 | ATGCGGGCGCGGGCGGCCTCGGCTGCGCCGAGAGCGGAGACACAGGCTCAAGATGGCAGATTCCGACTGAGGCTGGGGGGGCCGAGCTCGCGCGCCGCTTTCCCGTCCCCGTTGCCATGA | MRARAASAAPRAETQAQDGRFRLRLGGPSSRAAFPSPLP | 86.7 |
| cuORF038 | NM_003325 | ATGCGGCTGTGGTGGCGGCGGCGGCGGCCGAGCGCGGGTGGCGGCTGTGGCGGCGGAGGGGGGCGCGGGCCGGCGATGGCGCGGCGGCCCTGA | MRLWWRRRRPSAGGGCGGGGGRGPAMARRP | 65.8 |
| cuORF039 | NM_001127714 | ATGACTGGGCCATGGACACAGATGTCTCCAACCTTCAACCGTTTGCATAGCACACGGGGGACTCGTGGGGGCCACCTGCCACTGCCAGCTGAAACAATACAATGGCAATACTGA | MTGPWTQMSPTFNRLHSTRGTRGGHLPLPAETIQWQY | 97.1 |
| cuORF040 | NM_018411 | ATGGCGCAACCTACGGCCTCGGCCCAGAAGCTGGTGCGGCCGATCCGCGCCGTGTGCCGCATCCTGCAGATCCCGGAGTCCGACCCCTCCAACCTGCGGCCCTAG | MAQPTASAQKLVRPIRAVCRILQIPESDPSNLRP | 61.9 |
| cuORF041 | NM_007232 | ATGCGCTGGGCGCCCCCAGGGGAACCCGACCCGGCCAAGGGCCCGCAAAGACGAGGCTCCCGGGCCGGGGCCCCTCCCGGCCGCCCAGCTCTCGGCCGGCGCCCTGCCCCGCGTCCCGGAGCCGCGTGA | MRWAPPGEPDPAKGPQRRGSRAGAPPGRPALGRRPAPRPGAA | 89.7 |
| cuORF042 | NM_014920 | ATGTATTTGGGAGACAGTCACGTCCTATTGAATACCTTGTGCTGGTGCTGCCATCGAAAAATCTGGTTACACTCTGGGGAGGACTGCTACCACTGCAGAACTGAACCACTTCGGCCGTGA | MYLGDSHVLLNTLCWCCHRKIWLHSGEDCYHCRTEPLRP | 84.6 |
| cuORF043 | NM_001550 | ATGTATCGTTTTCGATCACAGCTCTTCACGGGGATTTCTGCTGCCGCCACCGCCCACTCTTACCCCCGCCGCTTCTCGACTCTGTTGTTAGCCGAAGACTCGCCTCTCAGCCGCCCGCCGCACAGACGCACGAGTAAAAAGTGCAGCTCCATCGGCTGA | MYRFRSQLFTGISAAATAHSYPRRFSTLLLAEDSPLSRPPHRRTSKKCSSIG | 78.4 |
| cuORF044 | NM_006390 | ATGGAGAGCGAGTTGTGGGGGGGAAAAAGGGAGGACAGGGGGCGCGGAGTCAGAGTGGCGCAGCAAGTGGCCGCAGGTGGCGACGGTGGCGGGGGGTGGGGTGTGAGGTAA | MESELWGGKREDRGRGVRVAQQVAAGGDGGGGWGVR | 71 |
| cuORF045 | NM_175913 | ATGGCACTGACTAGCCGACCCGGTGCTGACTTGGCGCCGGGGCTCTTTCAGCCTCTGCTGGCATCCTGCTTCCCCCCTCCTGGGGTGCAGTGGTGA | MALTSRPGADLAPGLFQPLLASCFPPPGVQW | 88.9 |
| cuORF046 | NM_152447 | ATGCGGGTGCTGAGCTATGGCAAAGGGCAGCGAAGTGACGAGCGAGACCCGCGTACGACTGTGAAAGCCACCTGGAGCCACCTTGCCGGGATTGTACCTGCAGGCAGAAAGTCTTCCTACGACCGTCTTTTCCCTTAG | MRVLSYGKGQRSDERDPRTTVKATWSHLAGIVPAGRKSSYDRLFP | 73.3 |
| cuORF047 | NM_152447 | ATGCTTTCACTCTGTCCGTCTTCTGCAGCAGCCAGGCAGAGTGCCGACTCCTTCACAGCCGTGAGGAACTCTTCAGGCTCCAGAAGCTCTTAA | MLSLCPSSAAARQSADSFTAVRNSSGSRSS | 60.6 |
| cuORF048 | NM_001031692 | ATGCAGCCCATTCTCTGGAGAACTTCCTCACACACCGCAGCAAAGAGAAGACTGAAAGACAAACCTGGGTGCAGCCAGAGAGGTCCAGATAGATGA | MQPILWRTSSHTAAKRRLKDKPGCSQRGPDR | 90 |
| cuORF049 | NM_032272 | ATGTCGCGGTGGCAGCCCGGATGGGCCGGCAGGGCCGGGAGTAACGGGACGTCGCCGCGGAGCTTCTTCCCCCGGATACAGTGCGGCCCGAGCGGAGGCCGCGGCGCCGCCCTCCGATCTTGA | MSRWQPGWAGRAGSNGTSPRSFFPRIQCGPSGGRGAALRS | 90.9 |
| cuORF050 | NM_018717 | ATGCGGTGTGGAGTGAGTGTGTGTCGCGCGTGTCCGCACTGGAGGCATATGCTTGTGTGTGTACATGGGGTGTGTTTTTCGGTATGTAGGGAGAAAATGCTTGCCAACCACCGGAAATCTCCTGGAATTTATTAG | MRCGVSVCRACPHWRHMLVCVHGVCFSVCREKMLANHRKSPGIY | 91.3 |
| cuORF051 | NM_002748 | ATGGCGGCGACTGCGGCAAAGCGAGAGCCTCGGAGACGCCGCTGCCGCCAGCACAGCCGGAGACCTGAGCCGACACTGGGGGCAGTCCGCGAGCCCCGCACTCTCTCGATGAGTCGGAGAAGTCCCGTTGTATCAGAGTAA | MAATAAKREPRRRRCRQHSRRPEPTLGAVREPRTLSMSRRSPVVSE | 93.3 |
| cuORF052 | NM_138396 | ATGGGGGGCGGCCCCGCCGCAGGAGGACCCGGGCGCGTAACCCCGGGGGCCCGGCCCGCTCCAGACCCAGACCCGCGGGGTGCGGCACCCTGA | MGGGPAAGGPGRVTPGARPAPDPDPRGAAP | 73.6 |
| cuORF053 | NM_020814 | ATGAAGGCCGGAGTCGCGACCGCGCCGGACGGCGGGCAGCAGCCAGAGGACGAGCCGGAGCAGCCCCCGCCCCGGAGACACCCGGACGCCGAGCAGCAGTCGCCTCCGCCGCCGCAGCAGCAGCTGCGGGCGCGGGCCGACCCGGAGCCGGAGGAGGAGGGCGACCGCGACCCAGAGGAGGAAGAGGAGGAGAAGGCGAGGCCACTCGCCCGCCCGGACCTGCCCTTCTCTCGGCTCTTGCCCCTCGACCGAAGGGACCTTTGA | MKAGVATAPDGGQQPEDEPEQPPPRRHPDAEQQSPPPPQQQLRARADPEPEEEGDRDPEEEEEEKARPLARPDLPFSRLLPLDRRDL | 83.3 |
| cuORF054 | NM_021647 | ATGGCGAGGGGTGCACGGCGGCCACCTGAGTGGCGCGGCGGTGTCAGGTTCTTGCTCAAGTACCAACTCTATGGACCCAGGACAGGTTTGTCCCATGACCTGCTGTGA | MARGARRPPEWRGGVRFLLKYQLYGPRTGLSHDLL | 87.1 |
| cuORF055 | NM_019008 | ATGGCCCCGTGGAGCCGAGAGGCGGTGCTGAGTCTCTATCGGGCTCTGTTGCGCCAGGGCCGACAGCTTCGCTACACTGATCGAGACTTCTACTTTGCCTCCATCCGCCGTGAATTCCGAAAAAATCAGAAGCTAGAGGACGCTGAGGCCCGGGAGAGGCAGCTGGAGAAGGGCCTGGTCTTTCTCAACGGCAAATTGGGGAGGATCATTTAG | MAPWSREAVLSLYRALLRQGRQLRYTDRDFYFASIRREFRKNQKLEDAEARERQLEKGLVFLNGKLGRII | 68.3 |
| cuORF056 | NM_170784 | ATGAGCCTTCGGAACTTGTGGAGAGACTACAAAGTTTTGGTTGTTATGGTCCCTTTAGTTGGGCTCATACATTTGGGGTGGTACAGAATCAAAAGCAGCCCTGTTTTCCAAATACCTAAAAACGACGACATTCCTGAGCAAGATAGTCTGGGACTTTCAAATCTTCAGAAGAGCCAAATCCAGGGGAAGTAG | MSLRNLWRDYKVLVVMVPLVGLIHLGWYRIKSSPVFQIPKNDDIPEQDSLGLSNLQKSQIQGK | 72 |

### (Continued)

| | | | | |
|---|---|---|---|---|
| cuORF057 | NM_170784 | ATGAAAAATACCAGTTGGATTAGAAAGAACTGGCTTCTTGTAGCTGGGATATCTTTCATAGGTGTCCATCTTGGAACATACTTTTTGCAGAGGTCTGCAAAGCAGTCTGTAAAATTTCAGTCTCAAAGCAAACAAAAGAGTATTGAAGAGTGA | MKNTSWIRKNWLLVAGISFIGVHLGTYFLQRSAKQSVKFQSQSKQKSIEE | 87.1 |
| cuORF058 | NM_005955 | ATGGTGCCGCTGGGGAGGGGAGAAGCTGCTGCTGCCGCCGTTGCCGGGAGCCGCGGAGACAAGTCATTACGTTTTCATTTCTCACAACTGGGCTGA | MVPLGRGEAAAAAVAGSRGDKSLRFHFSQLG | 83.9 |
| cuORF059 | NM_014751 | ATGGGCTTTGCCGTTTCCGTAGTGGGCACCAGTGGTGGCCTGATTGTCAGTCTTCTCCCGGCATTTTTAAGGCCAGGAGCCGAGCGCTGCTTGTAG | MGFAVSVVGTSGGLIVSLLPAFLRPGAERCL | 97 |
| cuORF060 | NM_006160 | ATGCCACACTCGCTCCGCGGTTCGCATGGCGCTCTGAAGACGCCGGCGCCCGCCGCCTTGAGGAGCCGCTGCCCCCGCTCCCTGAAGATGGGGGAACAATGA | MPHSLRGSHGALKTPAPAALRSRCPRSLKMGEQ | 84.8 |
| cuORF061 | NM_181829 | ATGCTGGCCGCTGGGGACCCGCGCAGCCCAGACCGTTCCCGGGCCGGCCAGCCGGCCACCATGGTGGCCCTGAGGCCTGTGCAGCAACTCCAGGGGGGCTAA | MLAAGDPRSPDRSRAGQPATMVALRPVQQLQGG | 66.7 |
| cuORF062 | NM_021724 | ATGCAAAGCAGAACTCCGTTGCCTCAACGTCCAACCCTTCTGCAGGGCTGCAGTCCGGCCACCCCAAGACCTTGCTGCAGGGTGCTTCGGATCCTGATCGTGAGTCGCGGGGTCCACTCCCCGCCCTTAGCCAGTGCCCAGGGGGCAACAGCGGCGATCGCAACCTCTAGTTTGAGTCAAGGTCCAGTTTGA | MQSRTPLPQRPTLLQGCSPATPRPCCRVLRILIVSRGVHSPPLASAQGATAAIATSSLSQGPV | 90.3 |
| cuORF063 | NM_003269 | ATGAACTTCGGGGACCCTTCGCCGCTGCGTGCGCGGCTCTCCCCGGAAACCCGGACCTGGCCGCCTCTTCCCTCGGAAGATTTCCCAGCAATCTAG | MNFGDPSPLRARLSPETRTWPPLPSEDFPAI | 64.8 |
| cuORF064 | NM_001004354 | ATGCGGCGGGGTCCCCGCGGCGGCGACCCCCGGCCCCGAGCGCCCGGAGCGCCCAGAGGCGGCGTGCGGGGCCCGGGGACGCCGCGCCCTCCATGCGCCGAGGCGCGCCCCGAGACAGCCGGGGGCCCGCGCCGCAGCCGCCGCCCGCGCTGA | MRRGPRGGDPRPRAPGAPRGGVRGPGTPRPPCAEARPETAGGPRRSRRPR | 66.7 |
| cuORF065 | NM_006179 | ATGTCAGGAAGGAGGGGGCCACCTGTGTCCTCCACAGGGGCCCCCCGAAGCCTGGGGACTCCCAGCCCCAGAGCTCCGAGGCGGAGGAGGTGCTGA | MSGRRGPPVSSTGAPRSLGTPSPRAPRRRRC | 69 |
| cuORF066 | NM_032961 | ATGTCAAGAAGAACATCCATCCGGAGAAATGAAGAGAATGAAAGTTTTAAGCTGCAGAGCCGTTCTGTGCTTTTCCGGCACAAAATTATATCGCTGATTTTAAGCCCTTTTGCATTTGCCAGCCGTTGA | MSRRTSIRRNEENESFKLQSRSVLFRHKIISLILSPFAFASR | 83 |
| cuORF067 | NM_203487 | ATGACTAACGACAAATCCGTGAATTTGTTCTCTGGAGTTGCTAATTTGCCAGCCCGAAGCAACACATTTTACAAGGAGGAAACATTTTGCTGCTTCTGA | MTNDKSVNLFSGVANLPARSNTFYKEETFCCF | 75.6 |
| cuORF068 | NM_022104 | ATGGCGGCAGCGGCGCTGGGGAGGGCGAGGCGGAGGCGGCAAAACGGGCGGTCGAGCAGAACGTGTAGCCGCGTCCCCTCCAGTCCGCTCCGGGCAGCTGCTGATGCAAGGAATCCCCTGGGCTCCCGTCCACTCCACTGCTGA | MAAAALGRARRRRQNGRSSRTCSRVPSSPLRAAADARNPLGSRPLHC | 87.9 |
| cuORF069 | NM_181523 | ATGGAGGACAGATGGACAGCCGTATGGCCAGTCACCTCTCCTCTTAAACCTTTGGAGAGTGGTCCTTTGTCCTCTGCTGGACACATAATAGGAATTCTAACACATTCTCTGAATTCACTTTTCATAAAAACGTAA | MEDRWTAVWPVTSPLKPLESGPLSSAGHIIGILTHSLNSLFIKT | 85.7 |
| cuORF070 | NM_138370 | ATGTGTCGCCCGCGAGGGGCCGGGGTCGGGGCCGCCGGGGCCATGCGCGCGGGCTGGGCAGGGGGCCGGCGGGGCGCAGAGCGGAGCCGCCTCGGAGCCTGA | MCRPRGAGVGAAGAMRAGWAGGRRGAERSRLGA | 73.9 |
| cuORF071 | NM_017761 | ATGTTGGCCTCAGCTCCTAGGCTGAACTCAGCAGATCGGCCCATGAAAACTTCTGTATTGAGACAAAGGAAGGGATCTGTCAGAAAGCAACACTTGTTATCTTGGGCTTGGCAGCAAGGAAGAGGACAGGTAGTGGAGATCCTGCAATCTGAAAAGCAGACTGAAAGGTGA | MLASAPRLNSADRPMKTSVLRQRKGSVRKQHLLSWAWQQGRGQVVEILQSEKQTER | 69.7 |
| cuORF072 | NM_001304332 | ATGGCCACATTCGGCTGCTGTCTGCATGTCTGGGGACACAGGGCACCCAGCCCCAGCCCTGAGAAGTTAAGCCAGGCCTTCTGGCTCAGTTCCACAGGGGCACTCCAGACCCCAGGCCCCTTTCAGAGCAAAAGCAACTGA | MATFGCCLHVWGHRAPSPSPEKLSQAFWLSSTGALQTPGPFQSKSN | 91.1 |
| cuORF073 | NM_181897 | ATGAGCAGGTATTGGGAGCCACAGACTGTATTTGGGAACCAGCTGCATGTCAGGACAAGCATAGATTGTGGTCTGTGGAGGCTGGGAGAGCACAATGCTTGA | MSRYWEPQTVFGNQLHVRTSIDCGLWRLGEHNA | 97.7 |
| cuORF074 | NM_000314 | ATGAGAGACGGCGGCGGCCGCGGCCCGGAGCCCCTCTCAGCGCCTGTGAGCAGCCGCGGGGGCAGCGCCCTCGGGGAGCCGGCCGGCCTGCGGCGGCGGCAGCGGCGGCGTTTCTCGCCTCCTCTTCGTCTTTTCTAA | MRDGGGRGPEPLSAPVSSRGGSALGEPAGLRRRQRRRFSPPLRLF | 94.9 |
| cuORF075 | NM_003463 | ATGATTAGGCCACAATCTTCAATGAGTAAACATATTCCTCAATTCTGTGGTGTTCTTGGTCACACATTTATGGAGTTTCTGAAGGGCAGTGGAGATTACTGCCAGGCACAGCACGACCTCTATGCAGACAAGTGA | MIRPQSSMSKHIPQFCGVLGHTFMEFLKGSGDYCQAQHDLYADK | 70.6 |
| cuORF076 | NM_080391 | ATGCCCAGTCCCCAGCTCGCCAGCGTTTTTCGTTGGAATATACGTTGCACATTTATGGCGATTCTGAGTGTGAGGGCAGACTTCTGCCAGGCTCAGCACAGCATTTTCGCTGACAAGTGA | MPSPQLASVFRWNIRCTFMAILSVRADFCQAQHSIFADK | 67.7 |
| cuORF077 | NM_015361 | ATGGGGTTAATTCCCTTTCGTAAGACTCTTACTTGCACCCACCCAGCCCCGCCGTCGCCCCGCCGCGCCGCGCTCCAACCGCCTCCTCCTCCTCAGTAA | MGLIPFRKTLTCTHPAPPSPRRAALQPPPPPQ | 91.2 |
| cuORF078 | NM_170672 | ATGCATTTCCCAATGATGCTAGCTGCCCTTAAAATTCTTTCAGATTTGGAACTGTATCTGTATGGAAACAACAGAGTTTTCTTGAAGTACAACTAA | MHFPMMLAALKILSDLELYLYGNNRVFLKYN | 65.9 |
| cuORF079 | NM_001100588 | ATGGAGTTTGAGGTCGTGAAACCTCCGCCGAGCTCTGCCCCGGCGAGACGAGGCGCCTCCGTCGCCAAGCCGCGCCGTCGCGGGGCTCCCGGGAGCGGCCCTTAG | MEFEVVKPPPSSAPARRGASVAKPRRRGAPGSGP | 67.7 |
| cuORF080 | NM_005612 | ATGTGCGGCTGCCGCGAGCTCGCGGCGCAGCAGCGGAGCGAGCGCCGCCGAGGCCCGGGGCCCCAGACCCTGGCGGCGGCTGCCGCAGCCGAGACGGCAGGGCGAGGCCCGGAGGCCTGA | MCGCRELAAQQRSERRRGPGPQTLAAAAAAETAGRGPEA | 62.2 |
| cuORF081 | NM_015178 | ATGCTGATCCGGAAGGGGCAGCGGCTGCAGTGCAGCGCGCGCGTCCGCTCCTGGGCCCACGTCCGGCCTGGGCTGCCCTGCCGGAGTTTCTGA | MLIRKGQRLQCSARVRSWAHVRPGLPCRSF | 77.4 |
| cuORF082 | NM_020724 | ATGAGAGAAGGCGAAGCCGGGCGAAGGAGCAGGACGCTCGGCAGCAGGACTCCTCCGCAAGCCCCGCGGGCCCTCTGTTCCCGCGCCCCCCGGGCGCCCAAGAAGCAAAATGCAACCACCGCGCTACAGCTGAGAGGGGAAGGGGCCAGCGCCATCCGCTCGCCCGGAATCCGGTCCCCGCGGCGGCTGCGCGTCCCGGCCGCGGCCCCTCCGGCCCGGGAGTGA | MREGEAGRRSRTLGSRTPPQAPRALCSRAPRAPKKQNATTALQLRGEGASAIRSPGIRSPRRLRVPAAAPPARE | 65.8 |
| cuORF083 | NM_001145845 | ATGTTGGGCGCCGCGCGGGGCTGGGGGCGCCCAGAAGACGTGCGAGTGTCCGCGGTCCTGCTGCTGTCTCCAGTACCCTCCGCATCCCCCAAGTGA | MLGAARGWGRPEDVRVSAVLLLSPVPSASPK | 71.4 |
| cuORF084 | NM_133368 | ATGTTGGGAATAAGCTCTGCAACTTTCTTTGGCATTCAGTTGTTAAAAACAAATAGGATGCAAATTCCTCAACTCCAGGTTATGAAAACAGTACTTGGAAAACTGAAAACTACCTAA | MLGISSATFFGIQLLKTNRMQIPQLQVMKTVLGKLKTT | 68.8 |
| cuORF085 | NM_153618 | ATGCTTTACGGAAGCGTCTTGGACAGGGTCTCCGCCAGGCGACAAGAGCTCGGTGCTGAGATGTGTTACGTTCTCATCTCCCCATCAATTATGGATGGAAACAAATAA | MLYGSVLDRVSARRQELGAEMCYVLISPSIMDGNK | 89.3 |
| cuORF086 | NM_015073 | ATGGTGGAGAACTGGACTCCACAGGCTCACAACCTTCCTGTCAGGTTTCAGGGCCCAGCATCCTTCATCCTGGGCCTGGCTGCCCTGAACAATGGCTGA | MVENWTPQAHNLPVRFQGPASFILGLAALNNG | 66.7 |

### (Continued)

| | | | | |
|---|---|---|---|---|
| cuORF087 | NM_080670 | ATGGCGGATGACAAGGATTCTCTGCCTAAGCTTAAGGACCTGGCATTTCTCAAGAACCAGCTGGAAAGCCTGCAGCGGCGTGTAGAAGACGAAGTCAACAGTGGAGTGGGCCAGGATGGCTCGCTGTTGTCCTCCCCGTTCCTCAAGGGATTCCTGGCTGGCTATGTGGTGGCCAAACTGAGGGCATCAGCAGTATTGGGCTTTGCTGTGGGCACCTGCACTGGCATCTATGCGGCTCAGGCATATGCTGTGCCCAACGTGGAGAAGACATTAAGGGACTATTTGCAGTTGCTACGCAAGGGGCCCGACTAG | MADDKDSLPKLKDLAFLKNQLESLQRRVEDEVNSGVGQDGSLLSSPFLKGFLAGYVVAKLRASAVLGFAVGTCTGIYAAQAYAVPNVEKTLRDYLQLLRKGPD | 94.4 |
| cuORF088 | NM_001286646 | ATGGGGATTACCAGCGGACCACCTGTTTGTATCACCTTCCCACCTCTCTGTAGCAAGAAAAATCACTTCAGCACCTCTTCAGTAATAATGATGAAACTGGTATTAACTGAATGGCAATTATGGATTTTTAACTCTAATTCACAGTAA | MGITSGPPVCITFPPLCSKKNHFSTSSVIMMKLVLTEWQLWIFNSNSQ | 76.9 |
| cuORF089 | NM_005585 | ATGTTGTCGCCCTGCGGCGCCCCTTCGACGACAGGCTGTGCGCGGTCTGCACGGCGCTCCGCGGCGGAGCTTCATGTGGGGCTGCGACCCGCGCAGCCGGCGCCTCGCTGA | MLSPCGAPSTTGCARSARRSAAELHVGLRPAQPAPR | 70 |
| cuORF090 | NM_017719 | ATGATTAGCGCTGGGTGCGGGGTTTCGGCGGCCGGGAGGGAGTTGTCGGCGCCGCGGCCGCTGCGGACGGACGCTCGCCTGCCGGCTGAGGAAAAAGAAGCAACTAACAAAACACTGTGA | MISAGCGVSAAGRELSAPRPLRTDARLPAEEKEATNKTL | 81.2 |
| cuORF091 | NM_080867 | ATGGCGATGGTGATGGCAGCAGTTACTCGCACAACCCCAGTTAAGCTGCGCTCCGGGAGATACATCCAGAAAGTGCCCAGAAGAAACTTCCTGCTGGAAAAAATGAAAAAGCAGTATTTATAA | MAMVMAAVTRTTPVKLRSGRYIQKVPRRNFLLEKMKKQYL | 66.7 |
| cuORF092 | NM_003107 | ATGCCAGTCTGTTGCATGCAGGCTTTTTGGCTTCCTACCTTGCAACAAAATAATTGCACCAACTCCTTAGTGCCGATTCCGCCCACAGAGAGTCCTGGAGCCACAGTCTTTTTTGCTTTGCATTGTAGGAGAGGGACTAAGTGCTAG | MPVCCMQAFWLPTLQQNNCTNSLVPIPPTESPGATVFFALHCRRGTKC | 78.2 |
| cuORF093 | NM_080862 | ATGGAGCTTGGGCCCCTGCCGCAGCCCGGTAGAGGCTGTGGAGGTCTACCGTCCGGAAGCCTGGTTCCCAGCCCCGTGGCCCATTCCTGGCTACGGGGAGTGGAGGCTCCCACGAGGTAG | MELGPLPQPGRGCGGLPSGSLVPSPVAHSWLRGVEAPTR | 80 |
| cuORF094 | NM_006927 | ATGAACCACAAGGACCCCGACTGCTCCAGACTGGATTATTTCAAGCCGCCAAAGAGGGGGACCCCTAGAGCTGGCAGCCAGCAATCCCAAGGGACTAGAGGGCTGCAATGGACTGACCTCCCCCTCACCAGGGTGGCAGGAGAGGCAGAGCCTCTGTGGCCTAGCTAG | MNHKDPDCSRLDYFKPPKRGTPRAGSQQSQGTRGLQWTDLPLTRVAGEAEPLWPS | 78.6 |
| cuORF095 | NM_020225 | ATGAAGTGTAATGCATTGTGGGACGTGTGTAAAATCGGAGCCTTCGCCGTGGGGGTGTGGGGGGGCGTGGGGAGGGCCGGACCCGCCGCTGGCGGTGTAGACGCCGACGAGGAGGGGCTGGGAAAATGTGCGCAGAGTCCGCCCGGGTCGTGCCCGCCGTAG | MKCNALWDVCKIGAFAVGVWGGVGRAGPAAGGVDADEEGLGKCAQSPPGSCPP | 75.8 |
| cuORF096 | NM_004865 | ATGGAGACTGGGGAGCGCGCCCGTCTCATCCTCATCCTTGTCCTCCAGCTTCTCCTTCGCATCCGACGCAACCGGCAGCAGCGCTGCCGCCGCGTCCTCAGCCACCGCTCCCTCTTCCCACGGATGTGA | METGERARLILILVLQLLLRIRRNRQQRCRRVLSHRSLFPRM | 80 |
| cuORF097 | NM_003239 | ATGAATGGAGTGCCTGAGAGACAAGTGTGTCCTGTACTGCCCCCACCTTTAGCTGGGCCAGCAACTGCCCGGCCCTGCTTCTCCCCACCTACTCACTGGTGA | MNGVPERQVCPVLPPPLAGPATARPCFSPPTHW | 75 |
| cuORF098 | NM_001204760 | ATGTGCAGACCGGATTCATCTTCTCGGAGCTGCGGCGGCGGCTTTGGGCTCAGGCGGCGGCGGCTCGCGCTCGGCCGCGGAGTCCTGGCAGCAGCGGGGACGCGGCGCGGGAGTCCGAGCTCTGGTGGCAGCTGA | MCRPDSSSRSCGGGFGLRRRRLALGRGVLAAAGTRRGSPSSGGS | 83.3 |
| cuORF099 | NM_014452 | ATGGGAAGTCGTTCCTTTGCTCTCTCGCGCCCAGTCCTCCTCCCTGGTTCTCCTCAGCCGCTGTCGGAGGAGAGCACCCGGAGACGCGGGCTGCAGTCGCGGCGGCTTCTCCCCGCCTGGGCGGCCGCGCCGCTGGGCAGGTGCTGA | MGSRSFALSRPVLLPGSPQPLSEESTRRRGLQSRRLLPAWAAAPLGRC | 86.5 |
| cuORF100 | NM_033502 | ATGATTGGCCGACCGCAGGAGAAGCCCCCAGAACCAGGCCCCCAACTCAGCCATCTGCGGAGGTCAAGGTGTGAGCGACGTCTCCTCACCACAGTGCTGTGTGGTCTATACCTCAGCCAGGGAGAGGATGTGAAACCCCCCGCCCTGCACATGAGTGGTACAGGCCAACAGGAACACCTGGCTCCAGCCACGTTCACAGACATGTCAGCCGTGGAGTAG | MIGRPQEKPPEPGPQLSHLRRSRCERRLLTTVLCGLYLSQGEDVKPPALHMSGTGQQEHLAPATFTDMSAVE | 90.9 |
| cuORF101 | NM_003355 | ATGACCATTAGGTGTTTCGTCTCCCACCCATTTTCTATGGAAAACCAAGGGGATCGGGCCATGATAGCCACTGGCAGCTTTGAAGAACGGGACACCTTTAGAGAAGCTTGA | MTIRCFVSHPFSMENQGDRAMIATGSFEERDTFREA | 75 |
| cuORF102 | NM_001039590 | ATGGTCTCTGCAAGATGGTTTATTCTTGAATTGGACCTTTTTAAGACTGACAAATGCTGGTACTTCATCTTCTATAAGTGGACTATAATTTCTTTTCTCAAGACAACTACATAA | MVSARWFILELDLFKTDKCWYFIFYKWTIISFLKTTT | 83.3 |
| cuORF103 | NM_198570 | ATGGACGGCGGCTCCCGGCTGGCGGCGGCGCGCCCCCGGGCTGTGAATGCGACTCGCCCCTCGGCCGCGCTCCCCGCCCGCCCGCCCGCCGGGACGTGGTAG | MDGGSRLAAARPRAVNATRPSAALPARPPAGTW | 80 |
| cuORF104 | NM_100264 | ATGGCTTCCCTCGCGCCCCACCGTCCTCTTCCGGAAGGCGGCTCCCTCCCTGCGCAGCCCGGAGCCCCTGAGATCAGCCTCGAGCAGGCGCCCGAGCGAGACTATCCCTAA | MASLAPHRPLPEGGSLPAQPGAPEISLEQAPERDYP | 97.1 |
| cuORF105 | NM_012477 | ATGGTGGCCTCAGCGAAGATGGGCCGGGCAGGGACCATGGCGGTGGCAGCAGAGGTGGCAGGGGCGGGGCGGCTGGCGGTAGAGGAGGCTGTGGTCCTCAGGGGGCTGTAG | MVASAKMGRAGTMAVAAEVAGAGRLAVEEAVVLRGL | 78.4 |
| cuORF106 | NM_023034 | ATGCGGCCTAGCTCGGTTCCGCCTCCTCCTCGCGCGGCCCCAGCGGCTGCCCGCACCCCAGCCCCACTCCGGGCCTCCGTGTCTCTCCTGTGA | MRPSSVPPPPRAAPAAARTPAPLRASVSLL | 97.9 |
| cuORF107 | NM_198581 | ATGTTGGTGAGGAGAAATCACCGTTACGGCCACTGTCCAAATCCCCGCGTCGCTGCACGCCGCCCGCCCGCTCCCACGCCACAGCCACCGGCGGCGAATAGAGACTAG | MLVRRNHRYGHCPNPRVAARRPPAPTPQPPAANRD | 86.1 |
| cuORF108 | NM_015457 | ATGAGCTGTATCTTGATCTGTCTGACTGTCCATGTTTTCCACCTGCAACCATTTGCATGTGTACAGCCTACTGTTTGTCTCCAGTTTTTAAACTGTACAAGTTGTGTTTCTTAA | MSCILICLTVHVFHLQPFACVQPTVCLQFLNCTSCVS | 84.8 |
| cuORF109 | NM_021224 | ATGAGACTCCCAAACAACTTCCACAACAATAACCCGAGCAGGAAGAGGAGAAAGAGAAAGAGGATAAGGAGGCGGTGGGGCTGGAGAACCCGAAGCACCTCCCGGCGCCGGGACGCTTCTTCTGTTCCTAATGTGAGAGGCTAG | MRLPNNFHNNNPSRKRRKRKRIRRRWGWRTRSTSRRRDASSVPNVRG | 66 |
| cuORF110 | NM_144631 | ATGAATGGAAGCGGCGGCGGCGGCGGGAGCGGCCTGAGCTGGGCGCCGGGGCCAGGGCCGGGGGCTGCCCAGGGCCCGCGCCGCTGCATGGGGGCGGCCCGCGGGCCCTGA | MNGSGGGGGSGLSWAPGPGPGAAQGPRRCMGAARGP | 100 |
| cuORF111 | NM_138447 | ATGGCGCTCCGGTCAATAAAATCGATAGCTGGAAGCTGCCTGTGTTCCAGGCAAAGGCGGTGCGGTAGCAGCGCCGCCATTTTCCCCGAAGGCATCTTCCGGTGCCTTTCACCCAAGTTCGGGCAGGAGTTTCCTGAATAA | MALRSIKSIAGSCLCSRQRRCGSSAAIFPEGIFRCLSPKFGQEFPE | 64.3 |

### Supplementary Table 2 | List of high-accumulation cuORFs that were previously reported.

| **High-accumulation cuORFs** | **Endogenous expression** | **Reference** |
|---|---|---|
| cuORF053 | - | - |
| cuORF087 | WB/MS | Rocha, A. L. *et al.* An inner mitochondrial membrane microprotein from the SLC35A4 upstream ORF regulates cellular metabolism. *J Mol Biol* 436, 168559 (2024). |
| cuORF002 | WB | Kitano, S. *et al.* Autonomous functionality of an upstream open reading frame in polycistronic mammalian mRNAs. 325571 Preprint at https://doi.org/10.1101/325571 (2018). |
| cuORF100 | - | - |
| cuORF082 | - | - |
| cuORF056 | WB | Akimoto, C. *et al*. Translational repression of the McKusick–Kaufman syndrome transcript by unique upstream open reading frames encoding mitochondrial proteins with alternative polyadenylation sites. *Biochimica et Biophysica Acta (BBA) - General Subjects* 1830, 2728–2738 (2013). |
| cuORF057 | WB | Akimoto, C. *et al*. Translational repression of the McKusick–Kaufman syndrome transcript by unique upstream open reading frames encoding mitochondrial proteins with alternative polyadenylation sites. *Biochimica et Biophysica Acta (BBA) - General Subjects* 1830, 2728–2738 (2013). |
| cuORF062 | - | - |

### Supplementary Table 3 | Synonymous substitution mutant sequences.

| cuORF_ID | label | sequence |
|---|---|---|
| cuORF001 | WT | ATGTGCCGGGCGGGGGGCCCGGGTTCGCCGAGCCGCAGGAGAGACACGCTGGGCCGACCCCAGAGAGGCGCTGGACAGGCTGGTGGTCCAGGCCGTGGTGCCTGCCAGGTGATGTGGGGCAAAGCCCCCCGCACAGGCCAC |
| | Mutant | ATGTGTAGAGCTGGAGGACCTGGATCTCCTTCTAGAAGAAGAGATACATTAGGAAGACCTCAAAGAGGAGCTGGACAAGCTGGAGGACCTGGAAGAGGAGCTTGTCAAGTTATGTGGGGAAAAGCTCCTAGAACAGGACAT |
| cuORF019 | WT | ATGCGGTGCGGGGCGGCCCGGTGCCCCCCCGCCCCGTCACGGCAGCCGCGGCGGCCGAGGGGACCGGGCCAGGGCCGGGGGCGGCGGCCCGAGCCGCGG |
| | Mutant | ATGAGATGTGGAGCTGCTAGATGTCCTCCTGCTCCTTCTAGACAACCTAGAAGACCTAGAGGACCTGGACAAGGAAGAGGAAGAAGACCTGAACCTAGA |
| cuORF028 | WT | ATGGTGGCAGTCACCTTGCCAGTGCAGCAGGCATGGATCAGACTGCAAGCTAGGGGCACGAGGCATCAGTTGGGAAGAGGGGACGCACAGCTAGGCTTGAGGCCTCTTCATCGGGATGTCCCCAGGCCCCCCAGCCCAGGCCCAGCA |
| | Mutant | ATGGTTGCTGTTACATTACCTGTTCAACAAGCTTGGATTAGATTACAAGCTAGAGGAACAAGACATCAATTAGGAAGAGGAGATGCTCAATTAGGATTAAGACCTTTACATAGAGATGTTCCTAGACCTCCTTCTCCTGGACCTGCT |
| cuORF038 | WT | ATGCGGCTGTGGTGGCGGCGGCGGCGGCCGAGCGCGGGTGGCGGCTGTGGCGGCGGAGGGGGGCGCGGGCCGGCGATGGCGCGGCGGCCC |
| | Mutant | ATGAGATTATGGTGGAGAAGAAGAAGACCTTCTGCTGGAGGAGGATGTGGAGGAGGAGGAGGAAGAGGACCTGCTATGGCTAGAAGACCT |
| cuORF041 | WT | ATGCGCTGGGCGCCCCCAGGGGAACCCGACCCGGCCAAGGGCCCGCAAAGACGAGGCTCCCGGGCCGGGGCCCCTCCCGGCCGCCCAGCTCTCGGCCGGCGCCCTGCCCCGCGTCCCGGAGCCGCG |
| | Mutant | ATGAGATGGGCTCCTCCTGGAGAACCTGATCCTGCTAAAGGACCTCAAAGAAGAGGATCTAGAGCTGGAGCTCCTCCTGGAAGACCTGCTTTAGGAAGAAGACCTGCTCCTAGACCTGGAGCTGCT |
| cuORF049 | WT | ATGTCGCGGTGGCAGCCCGGATGGGCCGGCAGGGCCGGGAGTAACGGGACGTCGCCGCGGAGCTTCTTCCCCCGGATACAGTGCGGCCCGAGCGGAGGCCGCGGCGCCGCCCTCCGATCT |
| | Mutant | ATGTCTAGATGGCAACCTGGATGGGCTGGAAGAGCTGGATCTAATGGAACATCTCCTAGATCTTTTTTTCCTAGAATTCAATGTGGACCTTCTGGAGGAAGAGGAGCTGCTTTAAGATCT |
| cuORF051 | WT | ATGGCGGCGACTGCGGCAAAGCGAGAGCCTCGGAGACGCCGCTGCCGCCAGCACAGCCGGAGACCTGAGCCGACACTGGGGGCAGTCCGCGAGCCCCGCACTCTCTCGATGAGTCGGAGAAGTCCCGTTGTATCAGAG |
| | Mutant | ATGGCTGCTACAGCTGCTAAAAGAGAACCTAGAAGAAGAAGATGTAGACAACATTCTAGAAGACCTGAACCTACATTAGGAGCTGTTAGAGAACCTAGAACATTATCTATGTCTAGAAGATCTCCTGTTGTTTCTGAA |
| cuORF068 | WT | ATGGCGGCAGCGGCGCTGGGGAGGGCGAGGCGGAGGCGGCAAAACGGGCGGTCGAGCAGAACGTGTAGCCGCGTCCCCTCCAGTCCGCTCCGGGCAGCTGCTGATGCAAGGAATCCCCTGGGCTCCCGTCCACTCCACTGC |
| | Mutant | ATGGCTGCTGCTGCTTTAGGAAGAGCTAGAAGAAGAAGACAAAATGGAAGATCTTCTAGAACATGTTCTAGAGTTCCTTCTTCTCCTTTAAGAGCTGCTGCTGATGCTAGAAATCCTTTAGGATCTAGACCTTTACATTGT |
| cuORF074 | WT | ATGAGAGACGGCGGCGGCCGCGGCCCGGAGCCCCTCTCAGCGCCTGTGAGCAGCCGCGGGGGCAGCGCCCTCGGGGAGCCGGCCGGCCTGCGGCGGCGGCAGCGGCGGCGTTTCTCGCCTCCTCTTCGTCTTTTC |
| | Mutant | ATGAGAGATGGAGGAGGAAGAGGACCTGAACCTTTATCTGCTCCTGTTTCTTCTAGAGGAGGATCTGCTTTAGGAGAACCTGCTGGATTAAGAAGAAGACAAAGAAGAAGATTTTCTCCTCCTTTAAGATTATTT |
| cuORF103 | WT | ATGGACGGCGGCTCCCGGCTGGCGGCGGCGCGCCCCCGGGCTGTGAATGCGACTCGCCCCTCGGCCGCGCTCCCCGCCCGCCCGCCCGCCGGGACGTGG |
| | Mutant | ATGGATGGAGGATCTAGATTAGCTGCTGCTAGACCTAGAGCTGTTAATGCTACAAGACCTTCTGCTGCTTTACCTGCTAGACCTCCTGCTGGAACATGG |

**Supplementary Table 4 | Comparison of genome-wide predictions of low-accumulation and translation-restricted uORFs. Numbers represent the counts of uORFs classified into each combination of predicted categories.**

| | **High translation-restricted probability** | **Low translation-restricted probability** |
|---|---|---|
| **High-accumulation** | 376 (0.55%) | 26,453 (38.9%) |
| **Low-accumulation** | 1,056 (1.55%) | 40,096 (59.0%) |

### Supplementary Table 5 | All primers used for plasmid construction.

| Plasmid | PCR template | Forward primer | Reverse primer |
|---|---|---|---|
| cuORF001 | cDNAs | GGAGACCCAAGCTTGCTAGCCACCATGTGCCGGGCGGGGGGC | CCTCCGGAGCCGCCGGATCCGTGGCCTGTGCGGGGGGC |
| cuORF002 | cDNAs | GGAGACCCAAGCTTGCTAGCCACCATGGACAGTCTGACAGAACAGAGAC | CCTCCGGAGCCGCCGGATCCGCGCGTTCCCAGGAAAAACAT |
| cuORF003 | cDNAs | GGAGACCCAAGCTTGCTAGCCACCATGGGACTCCAGTTGTCTCTGATCAC | CCTCCGGAGCCGCCGGATCCAAATAACAAGATTTTATGCCCTTTGGTGCA |
| cuORF004 | cDNAs | GGAGACCCAAGCTTGCTAGCCACCATGCTGCTGCTAGGGGTGG | CCTCCGGAGCCGCCGGATCCCAAAGTCTTTAGTGTCAGCCACGG |
| cuORF005 | cDNAs | GGAGACCCAAGCTTGCTAGCCACCATGCCCGAGTGCGACTGG | CCTCCGGAGCCGCCGGATCCCCGGCGCCCGTCCGAGTC |
| cuORF006 | cDNAs | GGAGACCCAAGCTTGCTAGCCACCATGCGAGATTTTCCTCTGGCAG | CCTCCGGAGCCGCCGGATCCCCGAGTTCCTCTCGGGTTGC |
| cuORF007 | cDNAs | GGAGACCCAAGCTTGCTAGCCACCATGGATGAAAAGAGGCTGCTGGA | CCTCCGGAGCCGCCGGATCCGTCACTCAGGCATTCCGTCTGC |
| cuORF008 | cDNAs | GGAGACCCAAGCTTGCTAGCCACCATGTCTTCTTGGGGCGCCA | CCTCCGGAGCCGCCGGATCCATATCGCAGTTTGAATTGTTCCGGC |
| cuORF009 | cDNAs | GGAGACCCAAGCTTGCTAGCCACCATGGAGTTTCTAGGGCGCGC | CCTCCGGAGCCGCCGGATCCCCCCTCCCGCTCCCGCTG |
| cuORF010 | cDNAs | GGAGACCCAAGCTTGCTAGCCACCATGCTGGCTGCTCTCTGGG | CCTCCGGAGCCGCCGGATCCGCTTGTGGCTTCCGACAGCT |
| cuORF011 | cDNAs | GGAGACCCAAGCTTGCTAGCCACCATGAACGGAATCCGGAGGCG | CCTCCGGAGCCGCCGGATCCCAAATGGCTGGAGGTGCTCG |
| cuORF012 | cDNAs | GGAGACCCAAGCTTGCTAGCCACCATGGCTGAAGACTACTGGGACG | CCTCCGGAGCCGCCGGATCCGAGCCGCGGGGCCGGCGC |
| cuORF013 | cDNAs | GGAGACCCAAGCTTGCTAGCCACCATGCGGCCCGTGGTCCTC | CCTCCGGAGCCGCCGGATCCGCAAACTAGCAATTCCAGCGCT |
| cuORF014 | cDNAs | GGAGACCCAAGCTTGCTAGCCACCATGAGCGATCGGGGCCCG | CCTCCGGAGCCGCCGGATCCGGGCCGGCCGGGCATGGC |
| cuORF015 | cDNAs | GGAGACCCAAGCTTGCTAGCCACCATGGGCCTGGCGCGGGCG | CCTCCGGAGCCGCCGGATCCGAGCGCGCGGCCCCCGCG |
| cuORF016 | cDNAs | GGAGACCCAAGCTTGCTAGCCACCATGAGGGAAAAAAACCCCGGAGC | CCTCCGGAGCCGCCGGATCCGCGAGACTCGACGAGGGTGA |
| cuORF017 | cDNAs | GGAGACCCAAGCTTGCTAGCCACCATGAGGCTCCGAGACAGGAC | CCTCCGGAGCCGCCGGATCCATATGGGCTCAGCTTCCACGAC |
| cuORF018 | cDNAs | GGAGACCCAAGCTTGCTAGCCACCATGTTAAAGATGAGCGGGTGGC | CCTCCGGAGCCGCCGGATCCGGTGTGGTGATGTATGAAGATACACTTCCT |
| cuORF019 | cDNAs | GGAGACCCAAGCTTGCTAGCCACCATGCGGTGCGGGGCGGCC | CCTCCGGAGCCGCCGGATCCCCGCGGCTCGGGCCGCCG |
| cuORF020 | cDNAs | GGAGACCCAAGCTTGCTAGCCACCATGTCGTTTTGTAGTCAGAGAAAACAGAGA | CCTCCGGAGCCGCCGGATCCGCTTCTCAGCACAGCGCAAG |
| cuORF021 | cDNAs | GGAGACCCAAGCTTGCTAGCCACCATGCGGCTGCTCCCTGCT | CCTCCGGAGCCGCCGGATCCGCCGTTCACGACCTCTGACAG |
| cuORF022 | cDNAs | GGAGACCCAAGCTTGCTAGCCACCATGGCGCGCTGTGCGGCG | CCTCCGGAGCCGCCGGATCCAGGGCTTGGGAAGGGACC |
| cuORF023 | cDNAs | GGAGACCCAAGCTTGCTAGCCACCATGCTTAGACTACTTAACATACAAACTGCT | CCTCCGGAGCCGCCGGATCCATAGAGATGGAGTGGAGTAGATATCCAGAA |
| cuORF024 | cDNAs | GGAGACCCAAGCTTGCTAGCCACCATGCAGCTGCCTGGAGAGAG | CCTCCGGAGCCGCCGGATCCAGGGCCCAGTCCTGGGCG |
| cuORF025 | cDNAs | GGAGACCCAAGCTTGCTAGCCACCATGCAAGAAAATTACAGTCCGCTGC | CCTCCGGAGCCGCCGGATCCACTTTCTCCCCTTCGGTCTTTCAT |
| cuORF026 | cDNAs | GGAGACCCAAGCTTGCTAGCCACCATGCCCTGTCCGGAGTGC | CCTCCGGAGCCGCCGGATCCCAGATAAGTCGAGGTCGATGCAGC |
| cuORF027 | cDNAs | GGAGACCCAAGCTTGCTAGCCACCATGCCTAGAAGGGCAGCGG | CCTCCGGAGCCGCCGGATCCGCAGCAACTCCTGGCCAGC |
| cuORF028 | cDNAs | GGAGACCCAAGCTTGCTAGCCACCATGGTGGCAGTCACCTTGC | CCTCCGGAGCCGCCGGATCCTGCTGGGCCTGGGCTGGG |
| cuORF029 | cDNAs | GGAGACCCAAGCTTGCTAGCCACCATGGCGGCGGCTCTGTCG | CCTCCGGAGCCGCCGGATCCATGGATCTCAATATGAACCTGCAGGC |
| cuORF030 | cDNAs | GGAGACCCAAGCTTGCTAGCCACCATGAAGAAGGGACAGAGCACGG | CCTCCGGAGCCGCCGGATCCGGCACTGCGCCCCGCGCC |
| cuORF031 | cDNAs | GGAGACCCAAGCTTGCTAGCCACCATGCCATCAGTCTGGGACGT | CCTCCGGAGCCGCCGGATCCGCTCACTGTCAAAGCCACCATCAG |
| cuORF032 | cDNAs | GGAGACCCAAGCTTGCTAGCCACCATGGTCAGGCGGCGGCGG | CCTCCGGAGCCGCCGGATCCGTGGCCGCGGCCCCGTAG |
| cuORF033 | cDNAs | GGAGACCCAAGCTTGCTAGCCACCATGTTGGCCAAATGTGCCCC | CCTCCGGAGCCGCCGGATCCGAGGTTTGTTAGCTTGGTCTCACAAAAGG |
| cuORF034 | cDNAs | GGAGACCCAAGCTTGCTAGCCACCCAGGAAAGCCTGGAAACACATTCAT | CCTCCGGAGCCGCCGGATCCCTTTTTTTCTCTTAATTCTTCATCCGGACT |
| cuORF035 | cDNAs | GGAGACCCAAGCTTGCTAGCCACCATGAGGAGGACCAACCATGAGC | CCTCCGGAGCCGCCGGATCCCAAAATTCCTCGCACCGACAACT |
| cuORF036 | cDNAs | GGAGACCCAAGCTTGCTAGCCACCATGTCCCCGGCGAATGGG | CCTCCGGAGCCGCCGGATCCGAGGCGGGCGGTGGCGGT |
| cuORF037 | cDNAs | GGAGACCCAAGCTTGCTAGCCACCATGCGGGCGCGGGCGGCC | CCTCCGGAGCCGCCGGATCCTGGCAACGGGGACGGGAAA |
| cuORF038 | cDNAs | GGAGACCCAAGCTTGCTAGCCACCATGCGGCTGTGGTGGCGG | CCTCCGGAGCCGCCGGATCCGGGCCGCCGCGCCATCGC |
| cuORF039 | cDNAs | GGAGACCCAAGCTTGCTAGCCACCATGACTGGGCCATGGACACAG | CCTCCGGAGCCGCCGGATCCGTATTGCCATTGTATTGTTTCAGCTGGC |
| cuORF040 | cDNAs | GGAGACCCAAGCTTGCTAGCCACCATGGCGCAACCTACGGCC | CCTCCGGAGCCGCCGGATCCGGGCCGCAGGTTGGAGGG |
| cuORF041 | cDNAs | GGAGACCCAAGCTTGCTAGCCACCATGCGCTGGGCGCCCCCA | CCTCCGGAGCCGCCGGATCCCGCGGCTCCGGGACGCGG |
| cuORF042 | cDNAs | GGAGACCCAAGCTTGCTAGCCACCATGTATTTGGGAGACAGTCACGTCC | CCTCCGGAGCCGCCGGATCCCGGCCGAAGTGGTTCAGT |

### (Continued)

| | | | |
|---|---|---|---|
| cuORF043 | cDNAs | GGAGACCCAAGCTTGCTAGCCACCATGTATCGTTTTCGATCACAGCTCTTCAC | CCTCCGGAGCCGCCGGATCCGCCGATGGAGCTGCACTT |
| cuORF044 | cDNAs | GGAGACCCAAGCTTGCTAGCCACCATGGAGAGCGAGTTGTGGGG | CCTCCGGAGCCGCCGGATCCCCTCACACCCCACCCCCC |
| cuORF045 | cDNAs | GGAGACCCAAGCTTGCTAGCCACCATGGCACTGACTAGCCGACC | CCTCCGGAGCCGCCGGATCCCCACTGCACCCCAGGAGG |
| cuORF046 | cDNAs | GGAGACCCAAGCTTGCTAGCCACCATGCGGGTGCTGAGCTATGG | CCTCCGGAGCCGCCGGATCCAGGGAAAAGACGGTCGTAGGAAGAC |
| cuORF047 | cDNAs | GGAGACCCAAGCTTGCTAGCCACCATGCTTTCACTCTGTCCGTCTTCT | CCTCCGGAGCCGCCGGATCCAGAGCTTCTGGAGCCTGAAGAGT |
| cuORF048 | cDNAs | GGAGACCCAAGCTTGCTAGCCACCATGCAGCCCATTCTCTGGAGAACT | CCTCCGGAGCCGCCGGATCCTCTATCTGGACCTCTCTGGCTGCAC |
| cuORF049 | cDNAs | GGAGACCCAAGCTTGCTAGCCACCATGTCGCGGTGGCAGCCC | CCTCCGGAGCCGCCGGATCCAGATCGGAGGGCGGCGCC |
| cuORF050 | cDNAs | GGAGACCCAAGCTTGCTAGCCACCATGCGGTGTGGAGTGAGTGT | CCTCCGGAGCCGCCGGATCCATAAATTCCAGGAGATTTCCGGTGGTTG |
| cuORF051 | cDNAs | GGAGACCCAAGCTTGCTAGCCACCATGGCGGCGACTGCGGCA | CCTCCGGAGCCGCCGGATCCCTCTGATACAACGGGACTTCTCCGAC |
| cuORF052 | cDNAs | GGAGACCCAAGCTTGCTAGCCACCATGGGGGGCGGCCCCGCC | CCTCCGGAGCCGCCGGATCCGGGTGCCGCACCCCGCGG |
| cuORF053 | cDNAs | GGAGACCCAAGCTTGCTAGCCACCATGAAGGCCGGAGTCGCG | CCTCCGGAGCCGCCGGATCCAAGGTCCCTTCGGTCGAGGG |
| cuORF054 | cDNAs | GGAGACCCAAGCTTGCTAGCCACCATGGCGAGGGGTGCACGG | CCTCCGGAGCCGCCGGATCCCAGCAGGTCATGGGACAAACCT |
| cuORF055 | cDNAs | GGAGACCCAAGCTTGCTAGCCACCATGGCCCCGTGGAGCCGA | CCTCCGGAGCCGCCGGATCCAATGATCCTCCCCAATTTGCCGT |
| cuORF056 | cDNAs | GGAGACCCAAGCTTGCTAGCCACCATGAGCCTTCGGAACTTGTGGAG | CCTCCGGAGCCGCCGGATCCCTTCCCCTGGATTTGGCTCTTCTG |
| cuORF057 | cDNAs | GGAGACCCAAGCTTGCTAGCCACCATGAAAAATACCAGTTGGATTAGAAAGAAC | CCTCCGGAGCCGCCGGATCCCTCTTCAATACTCTTTTGTTTGCTTTGAGA |
| cuORF058 | cDNAs | GGAGACCCAAGCTTGCTAGCCACCATGGTGCCGCTGGGGAGG | CCTCCGGAGCCGCCGGATCCGCCCAGTTGTGAGAAATGAAAACGT |
| cuORF059 | cDNAs | GGAGACCCAAGCTTGCTAGCCACCATGGGCTTTGCCGTTTCCGTAG | CCTCCGGAGCCGCCGGATCCCAAGCAGCGCTCGGCTCC |
| cuORF060 | cDNAs | GGAGACCCAAGCTTGCTAGCCACCATGCCACACTCGCTCCGC | CCTCCGGAGCCGCCGGATCCTTGTTCCCCCATCTTCAGGGAGC |
| cuORF061 | cDNAs | GGAGACCCAAGCTTGCTAGCCACCATGCTGGCCGCTGGGGAC | CCTCCGGAGCCGCCGGATCCGCCCCCCTGGAGTTGCTG |
| cuORF062 | cDNAs | GGAGACCCAAGCTTGCTAGCCACCATGCAAAGCAGAACTCCGTTGC | CCTCCGGAGCCGCCGGATCCAACTGGACCTTGACTCAAACTAGAGGTT |
| cuORF063 | cDNAs | GGAGACCCAAGCTTGCTAGCCACCATGAACTTCGGGGACCCTTCG | CCTCCGGAGCCGCCGGATCCGATTGCTGGGAAATCTTCCGAGGG |
| cuORF064 | cDNAs | GGAGACCCAAGCTTGCTAGCCACCATGCGGCGGGGTCCCCGC | CCTCCGGAGCCGCCGGATCCGCGCGGGCGGCGGCTGCG |
| cuORF065 | cDNAs | GGAGACCCAAGCTTGCTAGCCACCATGTCAGGAAGGAGGGGGCC | CCTCCGGAGCCGCCGGATCCGCACCTCCTCCGCCTCGG |
| cuORF066 | cDNAs | GGAGACCCAAGCTTGCTAGCCACCATGACTAACGACAAATCCGTGAATTTGTTC | CCTCCGGAGCCGCCGGATCCGAAGCAGCAAAATGTTTCCTCCTTGTAAAA |
| cuORF067 | cDNAs | GGAGACCCAAGCTTGCTAGCCACCATGTCAAGAAGAACATCCATCCGGAGA | CCTCCGGAGCCGCCGGATCCACGGCTGGCAAATGCAAAAGG |
| cuORF068 | cDNAs | GGAGACCCAAGCTTGCTAGCCACCATGGCGGCAGCGGCGCTG | CCTCCGGAGCCGCCGGATCCGCAGTGGAGTGGACGGGAGC |
| cuORF069 | cDNAs | GGAGACCCAAGCTTGCTAGCCACCATGGAGGACAGATGGACAGCCG | CCTCCGGAGCCGCCGGATCCCGTTTTTATGAAAAGTGAATTCAGAGAATG |
| cuORF070 | cDNAs | GGAGACCCAAGCTTGCTAGCCACCATGTGTCGCCCGCGAGGG | CCTCCGGAGCCGCCGGATCCGGCTCCGAGGCGGCTCCG |
| cuORF071 | cDNAs | GGAGACCCAAGCTTGCTAGCCACCATGTTGGCCTCAGCTCCTAGG | CCTCCGGAGCCGCCGGATCCCCTTTCAGTCTGCTTTTCAGATTGCAG |
| cuORF072 | cDNAs | GGAGACCCAAGCTTGCTAGCCACCATGGCCACATTCGGCTGC | CCTCCGGAGCCGCCGGATCCGTTGCTTTTGCTCTGAAAGGGGC |
| cuORF073 | cDNAs | GGAGACCCAAGCTTGCTAGCCACCATGAGCAGGTATTGGGAGCCAC | CCTCCGGAGCCGCCGGATCCAGCATTGTGCTCTCCCAGCC |
| cuORF074 | cDNAs | GGAGACCCAAGCTTGCTAGCCACCATGAGAGACGGCGGCGGC | CCTCCGGAGCCGCCGGATCCGAAAAGACGAAGAGGAGGCGAGAAAC |
| cuORF075 | cDNAs | GGAGACCCAAGCTTGCTAGCCACCATGATTAGGCCACAATCTTCAATGAGTAAA | CCTCCGGAGCCGCCGGATCCCTTGTCTGCATAGAGGTCGTGCT |
| cuORF076 | cDNAs | GGAGACCCAAGCTTGCTAGCCACCATGCCCAGTCCCCAGCTC | CCTCCGGAGCCGCCGGATCCCTTGTCAGCGAAAATGCTGTGCT |
| cuORF077 | cDNAs | GGAGACCCAAGCTTGCTAGCCACCATGGGGTTAATTCCCTTTCGTAAGACTC | CCTCCGGAGCCGCCGGATCCCTGAGGAGGAGGAGGCGGT |
| cuORF078 | cDNAs | GGAGACCCAAGCTTGCTAGCCACCATGCATTTCCCAATGATGCTAGCTG | CCTCCGGAGCCGCCGGATCCGTTGTACTTCAAGAAAACTCTGTTGTTTCC |
| cuORF079 | cDNAs | GGAGACCCAAGCTTGCTAGCCACCATGGAGTTTGAGGTCGTGAAACCT | CCTCCGGAGCCGCCGGATCCAGGGCCGCTCCCGGGAGC |
| cuORF080 | cDNAs | GGAGACCCAAGCTTGCTAGCCACCATGTGCGGCTGCCGCGAG | CCTCCGGAGCCGCCGGATCCGGCCTCCGGGCCTCGCCC |
| cuORF081 | cDNAs | GGAGACCCAAGCTTGCTAGCCACCATGCTGATCCGGAAGGGGC | CCTCCGGAGCCGCCGGATCCGAAACTCCGGCAGGGCAGC |
| cuORF082 | cDNAs | GGAGACCCAAGCTTGCTAGCCACCATGAGAGAAGGCGAAGCCGG | CCTCCGGAGCCGCCGGATCCCTCCCGGGCCGGAGGGGC |
| cuORF083 | cDNAs | GGAGACCCAAGCTTGCTAGCCACCATGTTGGGCGCCGCGCGG | CCTCCGGAGCCGCCGGATCCCTTGGGGGATGCGGAGGG |
| cuORF084 | cDNAs | GGAGACCCAAGCTTGCTAGCCACCATGTTGGGAATAAGCTCTGCAACTTTCT | CCTCCGGAGCCGCCGGATCCTTAGGTAGTTTTCAGTTTTCCAAGTACTGT |
| cuORF085 | cDNAs | GGAGACCCAAGCTTGCTAGCCACCATGCTTTACGGAAGCGTCTTGGAC | CCTCCGGAGCCGCCGGATCCTTTGTTTCCATCCATAATTGATGGGGAGAT |

### (Continued)

| | | | |
|---|---|---|---|
| cuORF086 | cDNAs | GGAGACCCAAGCTTGCTAGCCACCATGGTGGAGAACTGGACTCCAC | CCTCCGGAGCCGCCGGATCCGCCATTGTTCAGGGCAGCC |
| cuORF087 | cDNAs | GGAGACCCAAGCTTGCTAGCCACCATGGCGGATGACAAGGATTCTCT | CCTCCGGAGCCGCCGGATCCGTCGGGCCCCTTGCGTAG |
| cuORF088 | cDNAs | GGAGACCCAAGCTTGCTAGCCACCATGGGGATTACCAGCGGACC | CCTCCGGAGCCGCCGGATCCCTGTGAATTAGAGTTAAAAATCCATAATTG |
| cuORF089 | cDNAs | GGAGACCCAAGCTTGCTAGCCACCATGTTGTCGCCCTGCGGC | CCTCCGGAGCCGCCGGATCCGCGAGGCGCCGGCTGCGC |
| cuORF090 | cDNAs | GGAGACCCAAGCTTGCTAGCCACCATGATTAGCGCTGGGTGCG | CCTCCGGAGCCGCCGGATCCCAGTGTTTTGTTAGTTGCTTCTTTTTCCTC |
| cuORF091 | cDNAs | GGAGACCCAAGCTTGCTAGCCACCATGGCGATGGTGATGGCAG | CCTCCGGAGCCGCCGGATCCTAAATACTGCTTTTTCATTTTTTCCAGCAG |
| cuORF092 | cDNAs | GGAGACCCAAGCTTGCTAGCCACCATGCCAGTCTGTTGCATGCAG | CCTCCGGAGCCGCCGGATCCGCACTTAGTCCCTCTCCTACAATGC |
| cuORF093 | cDNAs | GGAGACCCAAGCTTGCTAGCCACCATGGAGCTTGGGCCCC | CCTCCGGAGCCGCCGGATCCCCTCGTGGGAGCCTCCAC |
| cuORF094 | cDNAs | GGAGACCCAAGCTTGCTAGCCACCATGAACCACAAGGACCCCGAC | CCTCCGGAGCCGCCGGATCCGCTAGGCCACAGAGGCTC |
| cuORF095 | cDNAs | GGAGACCCAAGCTTGCTAGCCACCATGAAGTGTAATGCATTGTGGGACG | CCTCCGGAGCCGCCGGATCCCGGCGGGCACGACCCGGG |
| cuORF096 | cDNAs | GGAGACCCAAGCTTGCTAGCCACCATGGAGACTGGGGAGCGC | CCTCCGGAGCCGCCGGATCCCATCCGTGGGAAGAGGGAGC |
| cuORF097 | cDNAs | GGAGACCCAAGCTTGCTAGCCACCATGAATGGAGTGCCTGAGAGACAAG | CCTCCGGAGCCGCCGGATCCCCAGTGAGTAGGTGGGGAGAAG |
| cuORF098 | cDNAs | GGAGACCCAAGCTTGCTAGCCACCATGTGCAGACCGGATTCATCTTCTC | CCTCCGGAGCCGCCGGATCCGCTGCCACCAGAGCTCGG |
| cuORF099 | cDNAs | GGAGACCCAAGCTTGCTAGCCACCATGGGAAGTCGTTCCTTTGCTCTC | CCTCCGGAGCCGCCGGATCCGCACCTGCCCAGCGGCGC |
| cuORF100 | cDNAs | GGAGACCCAAGCTTGCTAGCCACCATGATTGGCCGACCGCAG | CCTCCGGAGCCGCCGGATCCCTCCACGGCTGACATGTCTG |
| cuORF101 | cDNAs | GGAGACCCAAGCTTGCTAGCCACCATGACCATTAGGTGTTTCGTCTCCC | CCTCCGGAGCCGCCGGATCCAGCTTCTCTAAAGGTGTCCCGTTC |
| cuORF102 | cDNAs | GGAGACCCAAGCTTGCTAGCCACCATGGTCTCTGCAAGATGGTTTATTCTTG | CCTCCGGAGCCGCCGGATCCTGTAGTTGTCTTGAGAAAAGAAATTATAGTCC |
| cuORF103 | cDNAs | GGAGACCCAAGCTTGCTAGCCACCATGGACGGCGGCTCCCGGC | CCTCCGGAGCCGCCGGATCCCCACGTCCCGGCGGGCGGG |
| cuORF104 | cDNAs | GGAGACCCAAGCTTGCTAGCCACCATGGCTTCCCTCGCGCCC | CCTCCGGAGCCGCCGGATCCGGGATAGTCTCGCTCGGGCGC |
| cuORF105 | cDNAs | GGAGACCCAAGCTTGCTAGCCACCATGGTGGCCTCAGCGAAGATG | CCTCCGGAGCCGCCGGATCCCAGCCCCCTGAGGACCACAG |
| cuORF106 | cDNAs | GGAGACCCAAGCTTGCTAGCCACCATGCGGCCTAGCTCGGTTC | CCTCCGGAGCCGCCGGATCCCAGGAGAGACACGGAGGCCC |
| cuORF107 | cDNAs | GGAGACCCAAGCTTGCTAGCCACCATGTTGGTGAGGAGAAATCACCGTTAC | CCTCCGGAGCCGCCGGATCCGTCTCTATTCGCCGCCGGTG |
| cuORF108 | cDNAs | GGAGACCCAAGCTTGCTAGCCACCATGAGCTGTATCTTGATCTGTCTGACTGTC | CCTCCGGAGCCGCCGGATCCAGAAACACAACTTGTACAGTTTAAAAACTGGAG |
| cuORF109 | cDNAs | GGAGACCCAAGCTTGCTAGCCACCATGAGACTCCCAAACAACTTCCACAAC | CCTCCGGAGCCGCCGGATCCGCCTCTCACATTAGGAACAGAAGAAGC |
| cuORF110 | cDNAs | GGAGACCCAAGCTTGCTAGCCACCATGAATGGAAGCGGCGGCGGCG | CCTCCGGAGCCGCCGGATCCGGGCCCGCGGGCCGCCCC |
| cuORF111 | cDNAs | GGAGACCCAAGCTTGCTAGCCACCATGGCGCTCCGGTCAATAAAATC | CCTCCGGAGCCGCCGGATCCTTCAGGAAACTCCTGCCCGAAC |
| EGFP-cuORF001-FLAG | uORF-FLAG plasmid | GCATGGACGAGCTGTACAAGTGCCGGGCGGGGGG | CCTCCTCCGGAGCCGCCGGATCCGTGGCCTGTGCGGGGG |
| EGFP-cuORF013-FLAG | uORF-FLAG plasmid | GCATGGACGAGCTGTACAAGCGGCCCGTGGTCCTCC | CCTCCTCCGGAGCCGCCGGATCCGCAAACTAGCAATTCCAGCGCT |
| EGFP-cuORF017-FLAG | uORF-FLAG plasmid | GCATGGACGAGCTGTACAAGAGGCTCCGAGACAGGACGC | CCTCCTCCGGAGCCGCCGGATCCATATGGGCTCAGCTTCCACGAC |
| EGFP-cuORF019-FLAG | uORF-FLAG plasmid | GCATGGACGAGCTGTACAAGCGGTGCGGGGCGGC | CCTCCTCCGGAGCCGCCGGATCCCCGCGGCTCGGGCC |
| EGFP-cuORF022-FLAG | uORF-FLAG plasmid | GCATGGACGAGCTGTACAAGGCGCGCTGTGCGGC | CCTCCTCCGGAGCCGCCGGATCCAGGGCTTGGGAAGGGACCC |
| EGFP-cuORF027-FLAG | uORF-FLAG plasmid | GCATGGACGAGCTGTACAAGCCTAGAAGGGCAGCGGGTG | CCTCCTCCGGAGCCGCCGGATCCGCAGCAACTCCTGGCCAGC |
| EGFP-cuORF028-FLAG | uORF-FLAG plasmid | GCATGGACGAGCTGTACAAGGTGGCAGTCACCTTGCCAGT | CCTCCTCCGGAGCCGCCGGATCCTGCTGGGCCTGGGCTG |
| EGFP-cuORF029-FLAG | uORF-FLAG plasmid | GCATGGACGAGCTGTACAAGGCGGCGGCTCTGTCGG | CCTCCTCCGGAGCCGCCGGATCCATGGATCTCAATATGAACCTGCA |
| EGFP-cuORF030-FLAG | uORF-FLAG plasmid | GCATGGACGAGCTGTACAAGAAGAAGGGACAGAGCACGGAGC | CCTCCTCCGGAGCCGCCGGATCCGGCACTGCGCCCCGC |
| EGFP-cuORF032-FLAG | uORF-FLAG plasmid | GCATGGACGAGCTGTACAAGGTCAGGCGGCGGCGG | CCTCCTCCGGAGCCGCCGGATCCGTGGCCGCGGCCCC |
| EGFP-cuORF033-FLAG | uORF-FLAG plasmid | GCATGGACGAGCTGTACAAGTTGGCCAAATGTGCCCCATGT | CCTCCTCCGGAGCCGCCGGATCCGAGGTTTGTTAGCTTGGTCTCAC |
| EGFP-cuORF038-FLAG | uORF-FLAG plasmid | GCATGGACGAGCTGTACAAGCGGCTGTGGTGGCGGC | CCTCCTCCGGAGCCGCCGGATCCGGGCCGCCGCGCC |
| EGFP-cuORF047-FLAG | uORF-FLAG plasmid | GCATGGACGAGCTGTACAAGCTTTCACTCTGTCCGTCTTCTGC | CCTCCTCCGGAGCCGCCGGATCCAGAGCTTCTGGAGCCTGAAGAGT |
| EGFP-cuORF049-FLAG | uORF-FLAG plasmid | GCATGGACGAGCTGTACAAGTCGCGGTGGCAGCCC | CCTCCTCCGGAGCCGCCGGATCCAGATCGGAGGGCGGCGC |
| EGFP-cuORF051-FLAG | uORF-FLAG plasmid | GCATGGACGAGCTGTACAAGGCGGCGACTGCGGC | CCTCCTCCGGAGCCGCCGGATCCCTCTGATACAACGGGACTTCTCC |
| EGFP-cuORF068-FLAG | uORF-FLAG plasmid | GCATGGACGAGCTGTACAAGGCGGCAGCGGCGC | CCTCCTCCGGAGCCGCCGGATCCGCAGTGGAGTGGACGGGAG |
| EGFP-cuORF069-FLAG | uORF-FLAG plasmid | GCATGGACGAGCTGTACAAGGAGGACAGATGGACAGCCGTATG | CCTCCTCCGGAGCCGCCGGATCCCGTTTTTATGAAAAGTGAATTCA |

### (Continued)

| | | | |
|---|---|---|---|
| EGFP-cuORF070-FLAG | uORF-FLAG plasmid | GCATGGACGAGCTGTACAAGTGTCGCCCGCGAGGG | CCTCCTCCGGAGCCGCCGGATCCGGCTCCGAGGCGGCTC |
| EGFP-cuORF071-FLAG | uORF-FLAG plasmid | GCATGGACGAGCTGTACAAGTTGGCCTCAGCTCCTAGGCT | CCTCCTCCGGAGCCGCCGGATCCCCTTTCAGTCTGCTTTTCAGATT |
| EGFP-cuORF074-FLAG | uORF-FLAG plasmid | GCATGGACGAGCTGTACAAGAGAGACGGCGGCGGC | CCTCCTCCGGAGCCGCCGGATCCGAAAAGACGAAGAGGAGGCGAGA |
| EGFP-cuORF079-FLAG | uORF-FLAG plasmid | GCATGGACGAGCTGTACAAGGAGTTTGAGGTCGTGAAACCTCC | CCTCCTCCGGAGCCGCCGGATCCAGGGCCGCTCCCGGG |
| EGFP-cuORF084-FLAG | uORF-FLAG plasmid | GCATGGACGAGCTGTACAAGTTGGGAATAAGCTCTGCAACTTT | CCTCCTCCGGAGCCGCCGGATCCGGTAGTTTTCAGTTTTCCAAGTA |
| EGFP-cuORF088-FLAG | uORF-FLAG plasmid | GCATGGACGAGCTGTACAAGGGGATTACCAGCGGACCACC | CCTCCTCCGGAGCCGCCGGATCCCTGTGAATTAGAGTTAAAAATCC |
| EGFP-cuORF091-FLAG | uORF-FLAG plasmid | GCATGGACGAGCTGTACAAGGCGATGGTGATGGCAGCAG | CCTCCTCCGGAGCCGCCGGATCCTAAATACTGCTTTTTCATTTTTT |
| EGFP-cuORF092-FLAG | uORF-FLAG plasmid | GCATGGACGAGCTGTACAAGCCAGTCTGTTGCATGCAGGC | CCTCCTCCGGAGCCGCCGGATCCGCACTTAGTCCCTCTCCTACAAT |
| EGFP-cuORF096-FLAG | uORF-FLAG plasmid | GCATGGACGAGCTGTACAAGGAGACTGGGGAGCGCGC | CCTCCTCCGGAGCCGCCGGATCCCATCCGTGGGAAGAGGGAGC |
| EGFP-cuORF098-FLAG | uORF-FLAG plasmid | GCATGGACGAGCTGTACAAGTGCAGACCGGATTCATCTTCTCG | CCTCCTCCGGAGCCGCCGGATCCGCTGCCACCAGAGCTCGG |
| EGFP-cuORF102-FLAG | uORF-FLAG plasmid | GCATGGACGAGCTGTACAAGGTCTCTGCAAGATGGTTTATTCT | CCTCCTCCGGAGCCGCCGGATCCTGTAGTTGTCTTGAGAAAAGAAA |
| EGFP-cuORF103-FLAG | uORF-FLAG plasmid | GCATGGACGAGCTGTACAAGGACGGCGGCTCCCGG | CCTCCTCCGGAGCCGCCGGATCCCCACGTCCCGGCGGG |
| EGFP-cuORF109-FLAG | uORF-FLAG plasmid | GCATGGACGAGCTGTACAAGAGACTCCCAAACAACTTCCACAA | CCTCCTCCGGAGCCGCCGGATCCGCCTCTCACATTAGGAACAGAAG |

### Supplementary Table 6 | All DNA fragments used for plasmid construction.

| Plasmid | 1st fragment | 2nd fragment | 3rd fragment |
|---|---|---|---|
| cuORF041 | AAGCTTGCTAGCCACCATGCGGTGGGCCCCCCCCGGCGAGCCCGACCCCGCCAAGGGCCCCCA | CCGCCAAGGGCCCCCAGCGGCGGGGCAGCCGGGCCGGCGCCCCCCCCGGCCGGCCCGCCCTGG | CGGAGCCGCCGGATCCGGCGGCGCCGGGCCGGGGGGCGGGCCGCCGGCCCAGGGCGGGCCGGCC |
| cuORF103 | AAGCTTGCTAGCCACCATGGACGGCGGCAGCCGGCTGGCCGCCGCCCGGCCCCG | CCGCCGCCCGGCCCCGGGCCGTGAACGCCACCCGGCCCAGCGCCGCCCTGCCCG | CGGAGCCGCCGGATCCCCAGGTGCCGGCGGGGGGCCGGGCGGGCAGGGCGGCGCT |
| cuORF030 | AAGCTTGCTAGCCACCATGAAGAAGGGCCAGAGCACCGAGCGGCCCGGCCGGGCCG | CGGCCCGGCCGGGCCGCCGAGCCCGAGGCCGAGGCCCGGCTGGCCCCCGGGAGGG | CGGAGCCGCCGGATCCGGCGCTCCGGCCGGCGCCGCAGGGGCCCTCCCGGGGGGCCA |
| cuORF070 | AAGCTTGCTAGCCACCATGTGCCGGCCCCGGGGCGCCGGCGTGGGCGCCGCCGG | GCGTGGGCGCCGCCGGCGCCATGCGGGCCGGCTGGGCCGGCGGCCGGCGGGGCG | CGGAGCCGCCGGATCCGGCGCCCAGCCGGCTCCGCTCGGCGCCCCGCCGGCCGCC |
| cuORF079 | AAGCTTGCTAGCCACCATGGAGTTCGAGGTGGTGAAGCCCCCCCCCAGCAGCGCC | CCCCCCCAGCAGCGCCCCCGCCCGGCGGGGCGCCAGCGTGGCCAAGCCCCGGCGG | CGGAGCCGCCGGATCCGGGGCCGCTGCCGGGGGCGCCCCGCCGCCGGGGCTTGGCC |
| cuORF105 | AAGCTTGCTAGCCACCATGGTGGCCAGCGCCAAGATGGGCCGGGCCGGCACCATGGC | GGGCCGGCACCATGGCCGTGGCCGCCGAGGTGGCCGGCGCCGGCCGGCTGGCCGTGG | CGGAGCCGCCGGATCCCAGGCCCCGCAGCACCACGGCCTCCTCCACGGCCAGCCGGCC |
| cuORF051 | AAGCTTGCTAGCCACCATGGCCGCCACCGCCGCCAAGCGGGAGCCCCGGCGGCGGCGGTGCCGGCAG | GCGGCGGTGCCGGCAGCACAGCCGGCGGCCCGAGCCCACCCTGGGCGCCGTGCGGGAGCCCCGGACC | CGGAGCCGCCGGATCCCTCGCTCACCACGGGGCTCCGCCGGCTCATGCTCAGGGTCCGGGGCTCCCGC |
| cuORF001 | AAGCTTGCTAGCCACCATGTGCCGGGCCGGCGGCCCCGGCAGCCCCAGCCGGCGGCGGGACACCCTGG | CGGCGGGACACCCTGGGCCGGCCCCAGCGGGGCGCCGGCCAGGCCGGCGGCCCCGGCCGGGGCGCCTG | CGGAGCCGCCGGATCCGTGGCCGGTCCGGGGGGCCTTGCCCCACATCACCTGGCAGGCGCCCCGGCCGG |
| cuORF028 | AAGCTTGCTAGCCACCATGGTGGCCGTGACCCTGCCCGTGCAGCAGGCCTGGATCCGGCTGCAGGCCCGG | CCGGCTGCAGGCCCGGGGCACCCGGCACCAGCTGGGCCGGGGCGACGCCCAGCTGGGCCTGCGGCCCCTG | CGGAGCCGCCGGATCCGGCGGGGCCGGGGCTGGGGGGCCGGGGCACGTCCCGGTGCAGGGGCCGCAGGCCC |
| cuORF088 | AAGCTTGCTAGCCACCATGGGCATCACCAGCGGCCCCCCCGTGTGCATCACCTTCCCCCCCTGTGCAG | TCCCCCCCCTGTGCAGCAAGAAGAACCACTTCAGCACCAGCAGCGTGATCATGATGAAGCTGGTGCTGA | CGGAGCCGCCGGATCCCTGGCTGTTGCTGTTGAAGATCCACAGCTGCCACTCGGTCAGCACCAGCTTCAT |
| cuORF047 | AAGCTTGCTAGCCACCATGCTGAGCCTGTGCCCCAGCAGCGCCGCCGCCCG | GCAGCGCCGCCGCCCGGCAGAGCGCCGACAGCTTCACCGCCGTGCGGAACA | CGGAGCCGCCGGATCCGCTGCTCCGGCTGCCGCTGCTGTTCCGCACGGCGGT |
| cuORF038 | AAGCTTGCTAGCCACCATGCGGCTGTGGTGGCGGCGGCGGCGGCCCAGCGC | GGCGGCGGCCCAGCGCCGGCGGCGGCTGCGGCGGCGGCGGCGGCCGGGGCC | CGGAGCCGCCGGATCCGGGCCGCCGGGCCATGGCGGGGCCCCGGCCGCCGCC |
| cuORF049 | AAGCTTGCTAGCCACCATGAGCCGGTGGCAGCCCGGCTGGGCCGGCCGGGCCGGCAGCAAC | CCGGGCCGGCAGCAACGGCACCAGCCCCCGGAGCTTCTTCCCCCGGATCCAGTGCGGCCCC | CGGAGCCGCCGGATCCGCTCCGCAGGGCGGCGCCCCGGCCGCCGCTGGGGCCGCACTGGATC |
| cuORF074 | AAGCTTGCTAGCCACCATGCGGGACGGCGGCGGCCGGGGCCCCGAGCCCCTGAGCGCCCCCGTGAG | TGAGCGCCCCCGTGAGCAGCCGGGGCGGCAGCGCCCTGGGCGAGCCCGCCGGCCTGCGGCGGCGGC | CGGAGCCGCCGGATCCGAACAGCCGCAGGGGGGGGCTGAACCGCCGCCGCTGCCGCCGCCGCAGGCC |
| cuORF033 | AAGCTTGCTAGCCACCATGCTGGCCAAGTGCGCCCCCTGCAACAAGATGAAGCGGCGGGA | AGATGAAGCGGCGGGACAAGATGATGAGCTTCAGCCACATCGTGAAGCCCAAGACCAACG | CGGAGCCGCCGGATCCCAGGTTGGTCAGCTTGGTCTCGCAGAAGGCGTTGGTCTTGGGCTT |
| cuORF019 | AAGCTTGCTAGCCACCATGCGGTGCGGCGCCGCCCGGTGCCCCCCCGCCCCCAG | GCCCCCCCGCCCCCAGCCGGCAGCCCCGGCGGCCCCGGGGCCCCGGCCAGGGCC | CGGAGCCGCCGGATCCCCGGGGCTCGGGCCGCCGGCCCCGGCCCTGGCCGGGGCC |
| cuORF071 | AAGCTTGCTAGCCACCATGCTGGCCAGCGCCCCCCGGCTGAACAGCGCCGACCGGCCCATGAAGACCAGCGTGCTGC | AAGACCAGCGTGCTGCGGCAGCGGAAGGGCAGCGTGCGGAAGCAGCACCTGCTGAGCTGGGCCTGGCAGCAGGGCCG | CGGAGCCGCCGGATCCCCGCTCGGTCTGCTTCTCGCTCTGCAGGATCTCCACCACCTGGCCCCGGCCCTGCTGCCAGG |
| cuORF032 | AAGCTTGCTAGCCACCATGGTGCGGCGGCGGCGGCGGCGGCGGCGGCGGCGGCG | GGCGGCGGCGGCGGCGGCGGGGCGAGCCCAGCCCCCTGGGCTGCGGCGAGCTGC | CGGAGCCGCCGGATCCGTGGCCCCGGCCCCGCAGGCTCCGCAGCTCGCCGCAGCC |
| cuORF027 | AAGCTTGCTAGCCACCATGCCCCGGCGGGCCGCCGGCGGCCTGCGGGGCGCC | CGGCCTGCGGGGCGCCCTGCCCGCCGCCCTGCGGGGCGGCATCGAGGGCCTG | CGGAGCCGCCGGATCCGCAGCAGCTCCGGGCCAGCTTCAGGCCCTCGATGCCG |
| cuORF098 | AAGCTTGCTAGCCACCATGTGCCGGCCCGACAGCAGCAGCCGGAGCTGCGGCGGCGGCTTCGGCC | GGCGGCGGCTTCGGCCTGCGGCGGCGGCGGCTGGCCCTGGGCCGGGGCGTGCTGGCCGCCGCCGG | CGGAGCCGCCGGATCCGCTGCCGCCGCTGCTGGGGCTGCCCCGCCGGGTGCCGGCGGCGGCCAGCA |

### Supplementary Table 7 | RT-qPCR Primers used in this study.

| Target | Forward primer | Reverse primer |
|---|---|---|
| 18S ribosomal RNA | ATTAATCAAGAACGAAAGTCGGAGGT | TTTAAGTTTCAGCTTTGCAACCATACT |
| uORF plasmid 3'UTR | TAATAGAAGGGCGAATTCTGCAGATA TCC | TCGAGGCTGATCAGCGAGC |
